# Viperin immunity evolved across the tree of life through serial innovations on a conserved scaffold

**DOI:** 10.1101/2023.09.13.557418

**Authors:** H Shomar, H Georjon, Y Feng, B Olympio, F Tesson, J Cury, F Wu, A Bernheim

## Abstract

Evolutionary arms races between cells and viruses drive the rapid diversification of antiviral genes in diverse life forms. Recent discoveries have revealed the existence of shared immune genes between prokaryotes and eukaryotes, showing molecular and mechanistic similarities in their response to viruses. However, the underlying evolutionary dynamics that explain the conservation and adaptation of these antiviral genes remain mostly unexplored. Here, we show that viperins constitute a highly conserved family of immune genes across diverse prokaryotes and eukaryotes, and uncover mechanisms by which they diversified in eukaryotes. Our findings indicate that viperins are enriched in Asgard archaea and widely distributed in all major eukaryotic clades, suggesting their presence in the Last Eukaryotic Common Ancestor (LECA). We show that viperins maintain their immune function by producing antiviral nucleotide analogs. We demonstrate that eukaryotic viperins diversified through serial innovations on the viperin gene, such as the emergence and selection of substrate specificity towards pyrimidine nucleotides, and through partnerships with genes maintained through genetic linkage, notably with nucleotide kinases. These findings unveil biochemical and genomic transitions underlying the adaptation of immune genes shared by prokaryotes and eukaryotes. Our study paves the way for the understanding of the conservation of immunity across domains of life.

---

All cells, eukaryotic and prokaryotic, face the threat of viral infections^1,2^. The arms race between cells and viruses leads to a constant diversification of genes involved in antiviral defense which has been the focus of many evolutionary investigations on immunity^3–6^. Thus, many studies have primarily focused on clade-specific examinations of immune evolution^6–10^. Recent discoveries have shed light on the existence of immune systems shared between prokaryotes and eukaryotes^11–17^, predominantly humans, suggesting a previously overlooked aspect of immunity: conservation. Deciphering how these conserved immune genes evolve has proven challenging, as they often display structural and mechanistic similarities but little sequence homology^11–17^.

Viperins are enzymes involved in antiviral defense in bacteria and humans^18–20^. They display a uniquely high level of sequence homology for conserved immune genes with 42% identity between the human and the closest bacterial viperin. Viperins have been detected in animals, fungi, bacteria, and a few archaea^21–23^ suggesting an ancient origin^20^. These enzymes inhibit viral replication by producing modified nucleotides, converting nucleotide triphosphates (NTP) to their 3′-deoxy-3′,4′-didehydro (dhh) counterparts. Once integrated by RNA polymerases in the nascent RNA chain, ddh-nucleotides act as chain terminators and prevent further elongation of the RNA chain^18,20^. The conservation of the mechanism is such that the expression of the human viperin in *E. coli* protects the bacteria against T7 phage^20^. In this article, we investigate how this conserved family of immune genes evolved. We demonstrate that viperins are present and functionally conserved across archaeal and eukaryotic clades. We further uncover both genomic and biochemical transitions underlying the adaptation of viperins across domains of life.

## Viperins are an ancient family of immune genes present across the tree of life

Eukaryotes are suggested to have emerged 1.7-2.3 billion years ago from an ancestral archaeon most closely related to the present-day Asgard archaea^24^, yet the immune arsenal of the latter remains to be explored. To bridge this gap, we used DefenseFinder^25^ to perform a systematic detection of 152 antiviral systems in 496 Asgard archaeal genomes as well as 2289 other archaeal genomes (Fig 1a, Supplementary Fig. 1, Supplementary Table 1, Methods). Similar to bacteria, we show that archaea, including Asgard archaea, primarily rely on RM and CRISPR-Cas systems as their major antiviral defense mechanisms, while most other systems are encoded in less than 5% of genomes. However, we observe an enrichment of eukaryote-related immune genes such as pAgos and viperins in Asgard archaea compared to other archaea (Fig. 1b). Notably, this enrichment is most apparent for viperin genes, present in 6% of Asgard archaea, while only in 2% of other archaea, and in 0.5% of bacteria^25^. Within the Asgard archaea, viperins are one of the most abundant antiviral systems, and are found in over 10% of genomes across various sub-lineages that constitute the Heimdall group, which was proposed to be the closest to eukaryotes^26–28^ (Fig. 1c, inset). Hence, the immune arsenal of Asgard archaea shows widespread presence of systems that are not found in eukaryotes, such as RM and CRISPR-Cas, and other systems that are present in eukaryotes such as viperins. This suggests that CRISPR-Cas and RM have been likely lost early during the emergence of eukaryotes while viperins were transferred and persisted in eukaryotes.

**Figure 1.**
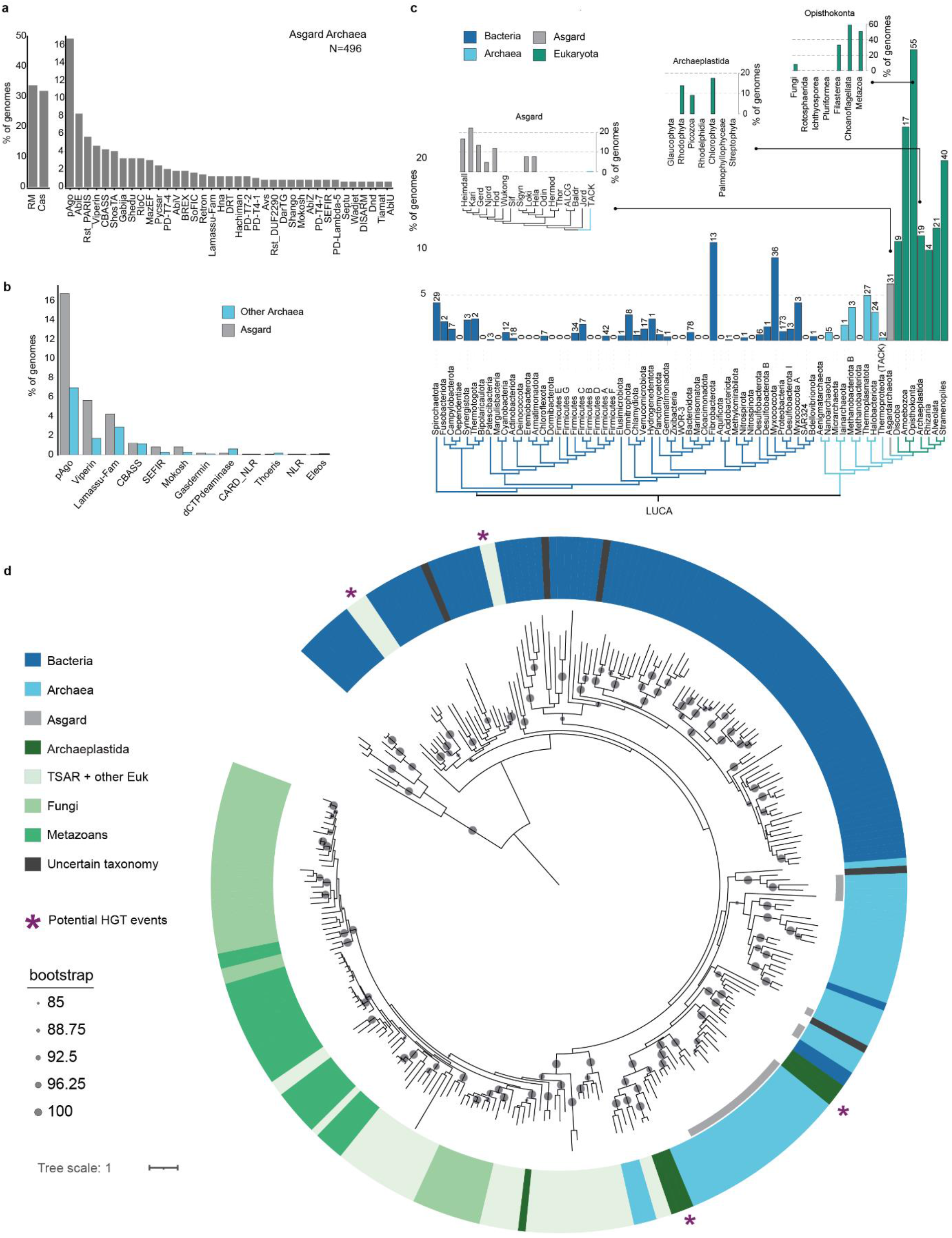
Viperins are an ancient family of immune genes present across domains of life. **a**. Detection of known prokaryotic defense systems across diverse Asgard archaea, ordered based on the percentage of analyzed genomes that encode them. **b**. Differential enrichment of eukaryote-related antiviral systems in Asgard archaea and other archaea. **c**. Distribution of viperins across bacteria, archaea and eukaryotes, where only lineages with more than 40 representatives are included. Vertical axis depicts the percentage of examined genomes encoding viperins. Numbers above bars indicate the number of viperins detected per lineage. Insets, heterogeneous distribution of viperins within Asgard archaea and selected eukaryotic supergroups. **d**. Phylogenetic tree of viperin rooted using MoaA as outgroup. Taxonomic classification is depicted by different colors. *indicate potential eukaryotic viperins acquired though HGT events from prokaryotes. Bootstrap values calculated through the UFBoot2+NNI approach are shown in gray circles.

Next, to explore the conservation of viperin-based immunity in eukaryotes, we sought to systematically examine their abundance and distribution across different lineages. We detected viperins in an expanded eukaryotic proteome database (see methods) and observed that all well-sampled eukaryotic clades (N>=40) retained viperin genes (Fig. 1c, Supplementary Table 2), ranging from 9% to 32% of organisms. Viperins thus appear to be generally more widespread in eukaryotes than in bacteria and archaea. However, similar to what was previously reported in prokaryotes, we observe a patchy distribution of viperins within eukaryotic supergroups. For example, viperins are found in some green algae (Chlorophyta) but are entirely absent in land plants (Streptophyta, N=35); within the Opisthokonta supergroup, viperins are 6 times more widespread in animals (Metazoa) than in fungi (Fig. 1c, inset). Thus, in both prokaryotes and eukaryotes, viperins are widespread but with a patchy distribution.

To further shed light on the evolutionary history of viperins, we built a phylogenetic tree using 264 representative viperins from taxonomically diverse eukaryotes and prokaryotes (Fig. 1d, Supplementary Fig. 2a, Supplementary Table 3). The overall topology of the phylogenetic tree, where archaea and bacteria form two independent branches with a deep root, suggests that viperins likely have an ancient origin in the Last Universal Common Ancestor (LUCA)^7^. Our analyses also show that the vast majority of eukaryotic viperins (EukVips) originated from a single ancient archaeal origin. Additionally, we observed four groups of 2-5 eukaryotic viperins within prokaryotic clades suggesting four independent events of horizontal gene transfer (HGT) (indicated with stars in Figure 1d). For example, Chlorophyta seemed to have acquired twice viperins through HGT, once directly from archaeal viperins and a second time through Cyanobacteria (Fig. 1d, Supplementary Fig. 2b-e). In five other cases, we only had one eukaryotic sequence embedded in prokaryotic viperins and could thus not rule out potential contamination (indicated in dark gray on Fig 1d). No HGT of viperins from eukaryotes to prokaryotes were observed. The major eukaryotic group is composed of a basal clade and a distal clade, with an apparent long phylogenetic distance (Supplementary Fig. 2e). The distal clade encompasses viperins from all eukaryotic supergroups, including all fungal and metazoan viperins. By contrast, the basal clade so far has only a small set of representatives and is only found in a specific clade named Diaphoretickes, which encompasses several supergroups including the highly sampled Archaeplastida (plants) and TSAR. However, the origin of the eukaryotic viperin branch is yet unresolved, as it robustly forms a sister clade to all archaeal viperins (Fig. 1d, Supplementary Fig. 2c-e, Methods). The diversity of Asgard archaea in nature is not yet fully explored, and their viperins are undersampled (Supplementary Fig. 2c). Given that Asgard viperins captured so far appear to have diverged early into multiple variants (Fig. 1d, Supplementary Fig. 2a,b,d), and viperins underwent frequent loss during evolution, it is plausible that an undiscovered or extinct Asgard archaeal lineage may carry a closer form to eukaryotic viperins. The phylogenetic patterns and taxonomic distributions observed in this study strongly indicates that viperins are an ancient family of genes that is present in the Last Eukaryotic Common Ancestor (LECA) and differentially lost during the subsequent events of evolution.

### The antiviral enzymatic activity of viperin is conserved in Asgard archaea and across diverse eukaryotes

Our genomics analysis revealed that viperins are a conserved and ancient gene family. We sought to establish whether the antiviral function of viperins is conserved across prokaryotes and eukaryotes. To assess the conservation of the enzymatic activity of viperins, we first analyzed the architecture of the active site across domains of life. We computed AlphaFold^29^ models of 92 viperin structures, encompassing homologs across the phylogenetic tree in Fig. 1d, and show a strong conservation of the active site architecture (Supplementary Fig. 3). Indeed, all viperins contain the previously reported partial (βα)_6_-barrel fold and C-terminal extension that result in an overall closed barrel structure similar to a (βα)_8_-TIM barrel, with conserved active site residues^23^(Fig. 2a). The structural conservation of the barrel fold led us to hypothesize that viperins remain catalytically active through their long-term evolution, while maintaining their antiviral function.

**Figure 2:**
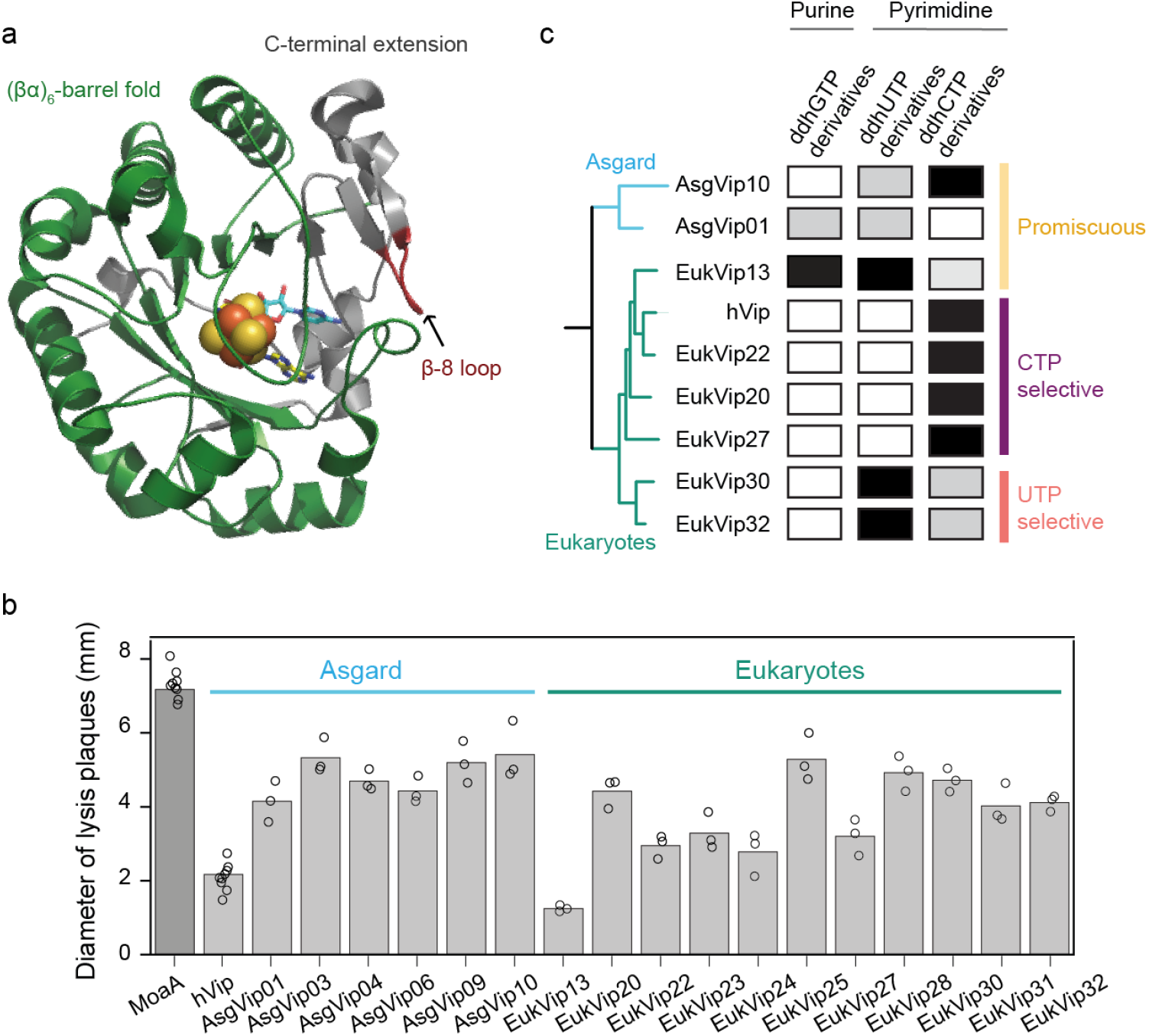
Asgard archaeal and eukaryotic viperins are antiviral and produce ddh-nucleotides. **a**. Crystal structure of mouse viperin^22^ (PDB ID: 6Q2P) illustrates the conserved active site architecture formed by a barrel fold, with conserved residues that ensure binding to the [4Fe-4S] cluster (shown as spheres), S-adenosylmethionine and CTP (shown as sticks). **b**. Diameter of phage T7 lysis plaques infecting *E. coli* expressing viperins or the control MoaA (dark gray). Bar graphs represent an average of three biological replicates, with individual data points overlaid. This graph only depicts viperin homologs that displayed a statistically significantly lower diameter compared to MoaA (one-sided t-test corrected by Bonferroni, p-value<0.01). Measurements for all viperins tested are in Supplementary Fig. 4. **c**. Production of ddh-nucleotide derivatives by diverse viperins. Black boxes depict detected compounds. Gray boxes depict detection of traces of ddh-derivatives. A ddh-nucleotide was defined as present, if at least 2 derivatives were found in all replicates above the limit of detection (LOD); compound traces are defined when only one derivative was found in all replicates above the LOD (see Supplementary Fig. 5). Phylogenetic tree on the left was computed with IQ-tree webserver (default settings) using the sequences of the viperins displayed.

Next, to test the immune function of previously unexplored viperins throughout the evolutionary spectrum of eukaryotes, we experimentally assessed the antiviral activity of viperins from diverse Asgard archaea and eukaryotes. To date, 26 viperins from diverse bacterial clades have been shown to confer defense against phages via heterologous expression in *E. coli*^20^, while the *in vivo* inhibition of viral replication has been confirmed for only 3 archaeal (from outside the Asgard clade^20^) and 2 eukaryotic (human and mouse^19,30^) viperins. Here, we cloned 35 viperin genes encoded by 10 Asgard archaea (AsgVips) and 25 eukaryotes (EukVips) for expression in *E. coli* (Supplementary Table 4). We evaluated the antiviral activity of these homologs by challenging viperin-expressing bacteria against phage T7, using the m*oaA* gene from *E. coli* as a negative control^20^. Remarkably, roughly half of the tested viperins exhibited clear anti-phage activity against T7 (6 of 10 AsgVips and 11 of 25 EukVips) (Fig. 2b, Supplementary Fig. 4), despite the large phylogenetic distance between *E. coli* and their native organisms. For instance, AsgVip01 from the recently discovered deep-sea *Candidatus Heimdallarchaeum aukensis*^28^, EukVip23 from the bird *Todus mexicanus*, and EukVip13 from jellyfish *Aurelia sp*. were all active in *E. coli*.

To further characterize these immune genes, we investigated the *in vivo* production of ddh-nucleotides by active eukaryotic and archaeal viperins. We used liquid chromatography-mass spectrometry (LC–MS) to analyze the metabolic content of *E. coli* cultures heterologously expressing viperins (Fig. 2c, Supplementary Fig. 5). First, we assessed whether our data aligns with previous results from *in vitro* assays using purified eukaryotic viperins. We show that the human viperin only produces ddhCTP, and EukVip30 (from *Trichoderma virens*) selectively produced ddhUTP, and some traces of ddhCTP in concordance with previous *in vitro* results^21,22^. Using this method, we examined the ddh-synthase activity of viperins from Asgard archaea and eukaryotes. We observed that the viperins AsgVip01 and AsgVip10 produce multiple ddh-nucleotides, showing that they are capable of promiscuous ddh-synthase activity, akin to certain bacterial viperins^20^. By contrast, we found that viperins from diverse eukaryotic organisms selectively produce one ddh-nucleotide variant, either ddhCTP or ddhUTP, suggesting substrate specificity. For example, EukVip20, EukVip22 and EukVip27 (from crustacea *Labidocera madurae*, fish *Alosa alosa* and foraminifera *Bolivina argentea*, Fig. 2c) synthesize exclusively ddhCTP, as observed for the human viperin. The fungal homologs, EukVip30 (from *Trichoderma virens*) and EukVip32 (from *Aspergillus coraciiforme*s), both catalyze mainly the production of ddhUTP, and some traces of ddhCTP. One exception is EukVip13 from *Aurelia sp*., which is capable of promiscuous ddh-synthase activity, as it produces ddh-nucleotides derived from CTP, UTP and GTP *in vivo* (Fig. 2c). This is the first evidence of a eukaryotic homolog that produces ddhGTP.

Overall, our results demonstrate that viperins conserved their function throughout the emergence and divergence of eukaryotes, via a common mechanism of ddhNTP synthesis, in accordance with previous in vitro studies^19,21,22^. Our data also suggest that there may be an evolutionary transition between archaea and eukaryotes, where viperins developed a preference for pyrimidine nucleotide substrates.

### Substrate specificity of viperins emerged and persisted in eukaryotes

Our results, combined with previous studies^21,22^, suggest that while the enzymatic activity of viperins is remarkably conserved, selectivity for pyrimidine nucleotide substrates may have specifically emerged in eukaryotes. To investigate a potential shift in substrate specificity and uncover evolutionary markers behind this trait, we sought to further characterize viperin residues involved in nucleotide binding. Previous analyses have revealed that the residues of the nucleobase binding site, located within the flexible β-8 loop of viperins, show significant variation between distant homologs^22^. Specific residues in the β-8 loop of previously crystalised viperins have been shown to coordinate substrate selectivity, and to discriminate between CTP and UTP substrates^21^ (Lys319 and Cys314 in *M. musculus*; Arg257 in *T. virens* respectively, Fig. 3a). We hypothesized that identifying potential stabilizing residues within the β-8 loop of distant homologs may elucidate the substrate selectivity of viperins across domains of life. Using 92 Alphafold models and a structure-guided alignment, we characterized the predicted nucleotide binding pocket of 1662 viperins (Supplementary Table 3). This structural analysis revealed three main pocket conformations with different stabilizing residues. We found numerous examples of viperins with the previously described CTP or a UTP selective pockets, comprising residues analogous to either Lys319/Cys314 from the mouse viperin, or Arg257 from the viperin of *T. virens* (Fig. 3a, Supplementary Fig. 6). Our LC-MS data confirmed that viperins with either a CTP (hVip, EukVip20, EukVip22, EukVip27) or a UTP (EukVip30, EukVip32) selective pocket only produce ddhCTP or ddhUTP *in vivo* (Fig. 2c). Finally, we identified a third type of binding pockets lacking any of these key residues. Most of these homologs have a highly conserved asparagine residue at the position of Cys314 in *M. musculus*, as exemplified in Fig. 3a. The structural models suggest that this conserved asparagine is the main stabilizing residue, which can theoretically interact with both UTP and CTP substrates. Moreover, the pocket of some of these homologs appears to provide a larger space, suggesting that it could potentially accommodate larger substrates such as purine nucleotides. Our results (Fig. 2c, AsgVip01, AsgVip10, EukVip13) show that viperins with this pocket can indeed use GTP as substrate, and that some are capable of promiscuous ddh-synthesis i*n vivo*. By determining the pocket conformation of previously studied viperins^20,21^ we show that most prokaryotic homologues have a pocket with a conserved asparagine residue, and that they can also synthesize ddhGTP and display promiscuous ddh-synthase activity. In addition, homologues that we determined to have this type of pocket but produced only one type of ddh-nucleotide *in vivo*^20^ (i.e. pVip8, pVip62 and pVip56), were previously shown to produce multiple ddh-nucleotides *in vitro* (including ddhATP)^31^. Altogether, these observations indicate a higher degree of substrate promiscuity among viperins with a stabilizing asparagine within the β-8 loop.

**Figure 3:**
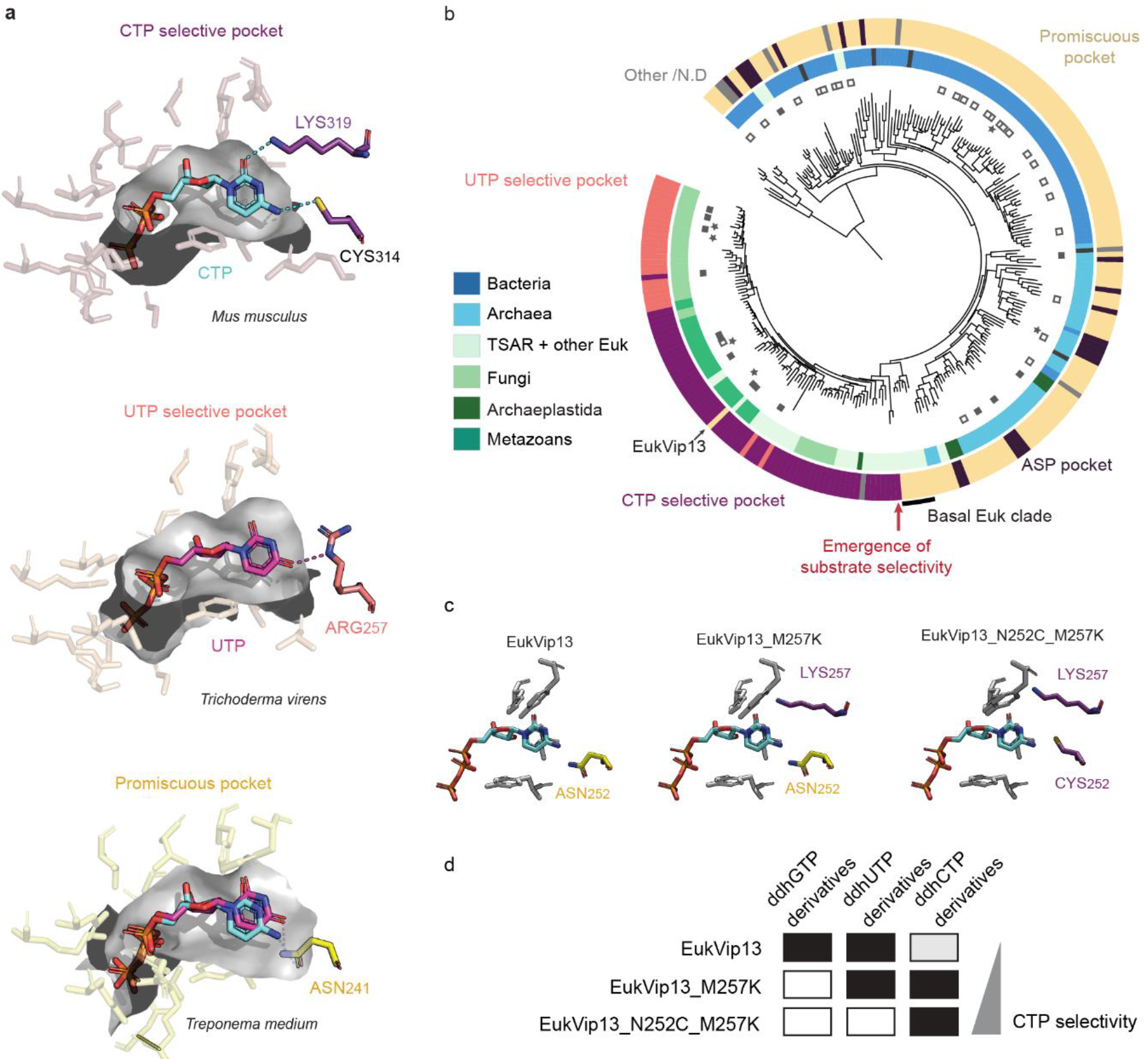
Viperin substrate specificity emerged in eukaryotes through serial innovations. **a**. Magnified cutaway of representative nucleotide binding pockets in viperin structures. Residues within the β-8 loop that interact with the nucleobase are highlighted. Residues Lys319 and Cys314 of the mouse viperin (PDB ID: 6Q2P), stabilize the selective binding of CTP, by respectively interacting with the C2 exocyclic oxygen and the C4 exocyclic amine of the cytidine base. In the viperin from *T. virens* (PDB ID: 7N7I), Arg257 selectively coordinates the uridine base, forming a hydrogen bond with its C4 exocyclic oxygen. The predicted promiscuous pockets are characterized by a highly conserved asparagine residue - Asn241 in viperin from *Treponema medium* (AlphaFold model). **b**. Phylogenetic tree of viperin (from Fig. 1d) annotated with corresponding nucleotide binding pockets (colored outer ring). Variants of the CTP selective (purple), UTP selective (pink) and ASP pockets (dark purple) are described in Supplementary Fig. 6 and Supplementary Fig. 7. Squares indicate viperins characterized *in vivo* in this study (black squares) and in previous studies (empty squares). Stars indicate viperins previously characterized *in vitro*. **c**. AlphaFold models of the nucleotide binding pockets of EukVip13 and mutated variants. Residues predicted to interact with the nucleobase are highlighted. **d**. Production of ddh-nucleotide derivatives by EukVip13 and mutated variants. Black boxes depict detected compounds. Gray boxes depict detection of traces of ddh-derivatives.

To examine how substrate selectivity emerged during evolution, we mapped the results of our structural analysis on the phylogenetic tree of viperins (Fig. 3b). We observed that the vast majority of viperins from bacteria and archaea have the identified promiscuous pocket, while both selective pockets are exclusively found in eukaryotes. The above identified EukVip13 appears as a singleton with substrate promiscuity in the distal eukaryotic branch, indicating that its asparagine residue emerged recently during its evolution. Strikingly, the promiscuous pocket is instead found across the entire basal clade of eukaryotic viperins, suggesting that viperins likely displayed promiscuous ddh-synthase activity in earliest eukaryotes. The CTP selective pocket is present only in the distal clade that encompasses viperins from all major eukaryotic supergroups, suggesting the emergence of substrate specificity within this eukaryotic branch followed by selection of this pocket. Whereas CTP is the preferred substrate for most eukaryotic viperins, the UTP selective pocket emerged as a unique innovation in a specific clade of fungal viperins. In addition, we report viperins with rare pocket conformations, referred to as ASP pockets, characterized by a conserved aspartic acid residue at the position of Cys314, and diverse residues at the position of Lys319 of the mouse viperin (Supplementary Fig. 7). While the ASP pocket is found sporadically among viperins in prokaryotes and eukaryotes, the eukaryotic viperins harboring these pockets are only found outside the main eukaryotic clade. This indicates that eukaryotic viperins observed with these binding sites were exclusively acquired through HGT from prokaryotes. Altogether, these observations allow us to track the emergence of substrate selectivity to early eukaryotes, suggesting a strong evolutionary advantage in producing pyrimidine ddh-nucleotides (in particular ddhCTP) as an innate immune response against eukaryotic viruses.

Our hypothesis about the emergence of the substrate specificity in eukaryotes is built on our structural analysis, and the assumption that the identified residues are the primary determinants of selective nucleotide binding. This would imply that mutations of these key residues in the β-8 loop would be sufficient to modify viperin substrate selectivity. To validate this hypothesis, we aimed to modify the substrate specificity of a promiscuous viperin, to make it selective for CTP. The protein EukVip13 is a unique homolog from the main eukaryotic clade with a promiscuous pocket (Fig. 3b). We generated variants of EukVip13 with mutations in key residues hypothesized to determine substrate selectivity (EukVip13_M257K and EukVip13_N252C_M257K) and used Alphafold^29^ models to visualize the hypothesized pocket conformations (Fig. 3c). By analyzing the metabolic content of *E. coli* expressing EukVip13 and mutated variants, we observe that EukVip13_M257K acquired substrate selectivity to pyrimidine substrates, while EukVip13_N252C_M257K displays high-level selectivity for CTP as expected. This demonstrated that as few as 2 point mutations are sufficient to transition from a promiscuous to a selective pocket. Hence, the extreme rarity of promiscuous viperins in eukaryotes suggests a strong selective pressure to maintain their substrate specificity towards CTP, while a small branch of eukaryotes (fungi) evolved selectivity for UTP. Overall, our findings show the exceptional conservation of the viperin family and how in this conserved framework, adaptation to eukaryotes happened through emergence and selection of substrate specificity towards pyrimidine nucleotides.

### Eukaryotic viperins innovated through functional genetic linkage

Following the observation that substrate selectivity emerged in eukaryotic viperins, we sought to explore additional adaptation mechanisms of viperins in eukaryotes. First, we evaluated whether specific regions of viperins showed more variability than others across domains of life. The sequence alignment of distant viperins shows significant variation in the N-terminal region that precedes the conserved β-barrel as previously reported ^23,32^(Figure 4.a). We observe a clear distinction between eukaryotic and prokaryotic viperins regarding the presence of a variable N-terminal tail (N-tails), with 83% of the eukaryotic viperins we analyzed having this feature, compared to only 10% of prokaryotic viperins (Supplementary Table 3). These N-tails are often predicted to form a transmembrane alpha-helix, but only in eukaryotes (37% of eukaryotic N-tails). Interestingly we observe that such transmembrane N-tails seem to have emerged early in the evolution of eukaryotes, as they are present in viperins from very diverse eukaryotic organisms, ranging from TSAR, metazoans and fungi (Supplementary Fig. 8). The presence of transmembrane N-tails has been previously reported in diverse mammalian viperins^23,32^ and they have been demonstrated to be crucial for antiviral activity in eukaryotes, as they target the protein to the endoplasmic reticulum (ER), where many viral replication processes occur^33^. The precise subcellular localization and role of all the detected N-tails is unknown. Overall, N-tails constitute a eukaryotic trait that could reflect a key evolutionary adaptation to the emergence of eukaryotic subcellular compartments and their role in viral replication.

**Figure 4:**
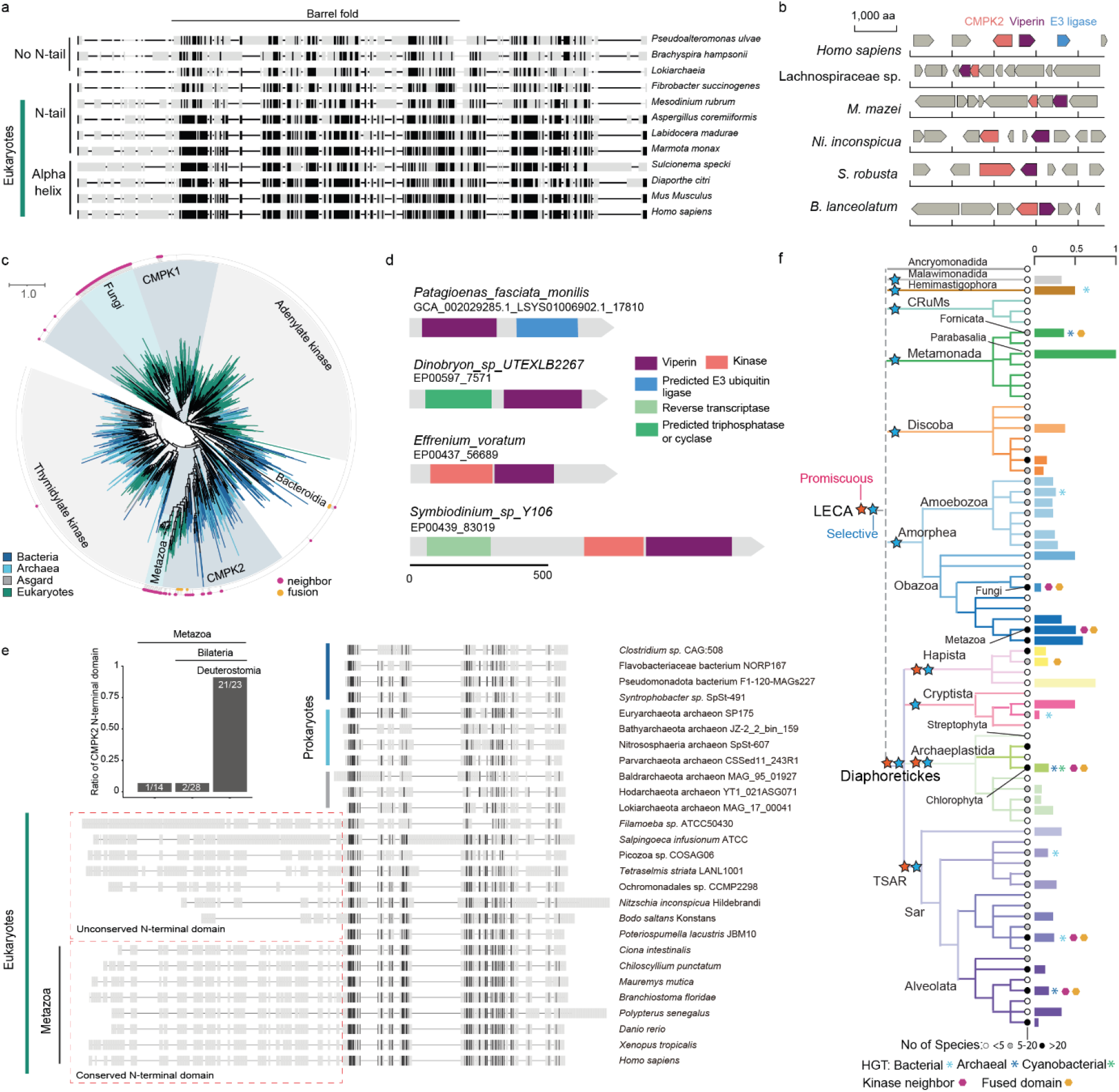
Eukaryotic viperins adapted through functional genetic linkage. **a**. Multiple sequence alignment of diverse viperins. Black indicates amino acids conserved in at least 50% of the sequences. **b**. Genomic depiction of viperins and their associated kinases in diverse organisms **c**. Phylogenetic tree of CMPK2 and their related kinases. Outer rim circles indicate the presence of viperin associated kinases. The blue shades highlight various representative forms of viperin-associated kinases. **d**. Examples of fusion between viperin and various functional domains. **e**. Sequence alignment of CMPK2 kinases indicates the emergence of an N-terminal extension in eukaryotes, and the fixation of the human CMPK2-like NTD in Metazoa. Black indicates amino acids conserved in at least 50% of the sequences. f. Distribution of viperins across the eukaryotes. Blue and red stars respectively show the selective and promiscuous versions of eukaryotic viperins. Bar plot shows the fraction of genomes encoding viperins.

Beyond the viperin gene itself, we hypothesized that viperins could adapt by functional association with other genes. Indeed, it was previously demonstrated that the human viperin works in concert with the neighboring kinase CMPK2, which enhances the conversion of CMP to CTP, hence providing the substrate for antiviral activity^34^. Strikingly, many prokaryotic viperins are found next to a kinase gene (Figure 4.b)^20^. In prokaryotes, genetic linkage (in the form of gene fusions, operons or islands) frequently indicates functional association. However, such functional genetic linkage is less prevalent in eukaryotes. Thus, we aimed to explore whether the viperin-kinase genetic linkage was conserved across the tree of life. We examined the gene neighborhoods of all viperins detected in this study. We found 484 nucleotide kinase genes in the vicinity of viperins (less than 3 genes away). 12% of the bacterial viperin genes, 5% of the archaeal viperin genes, and 70% of fungal and metazoan viperins genes encode an associated kinase (Figure 4b, Supplementary Fig. 9, Supplementary Table 5). We then evaluated whether these kinases belonged to the same family or were acquired through distinct events. CMPK2 is part of the large superfamily of nucleoside monophosphate kinases (NMPKs), that encompasses CMPK1, adenylate kinases and thymidylate kinases. We built a phylogenetic tree using representatives of this superfamily and the detected viperin-associated kinases (Methods, Figure 4.c). Notably, we observe that most viperin-associated kinases are evolutionarily related to the kinase domain of the human CMPK2 (Fig. 4c, Supplementary Fig. 9, Supplementary Fig. 10), showing that the viperin-kinase association was vertically inherited and maintained in eukaryotes. Interestingly, certain NMPKs associated with viperins from fungi (CMPK1) and two small clades of bacteria (unclassified) have distinct evolutionary origins, suggesting multiple lines of convergent evolution. This also coincides with the divergent, UTP-specific substrate preference in fungal viperins, implicating a potential role of CMPK1 in supporting the UTP-specific viperin functions.

Gene fusion is an extreme case of genetic linkage. Our analysis revealed 45 examples of eukaryotic viperins fused to additional protein domains (Supplementary Table 6). The most prevalent fusion we observe corresponds to nucleotide kinases (18 viperins), which frequently contain a CMPK2 domain (Figure 4.c). Beyond nucleotide kinases, we found instances of viperins fused to diverse additional domains (Figure 4.d), including a predicted E3-ubiquitin ligase domain (5 viperins), a predicted triphosphatase or cyclase domain (4 viperins), and even a reverse-transcriptase domain (1 viperin). Interestingly, it has been shown that the human viperin can activate the protein ubiquitination machinery by interacting with the E3 ubiquitin ligase ^35,36^, which is also found in the genomic vicinity in humans (Figure 4.b). The genomic fusion of viperin with a ubiquitin ligase further supports the hypothesis that viperin could work in concert with E3 ubiquitin ligase in different organisms. This also suggests that studying domains fused to the viperin gene could reveal additional proteins that work in concert with eukaryotic viperins.

To further understand the functional association of viperins and the CMPK2 family, we surveyed the evolution of CMPK2 across kingdoms. It was recently shown that the human CMPK2 contains a N-terminal domain (NTD) that displays a mitochondria-targeting sequence and antiviral activity, independent of viperin^37^. To study the emergence of this NTD, we aligned representative sequences of CMPK2 from diverse prokaryotic and eukaryotic organisms. Interestingly, we found that an N-terminal extension is only found in a subset of eukaryotes (Fig. 4e). The NTD from the human CMPK2 was found in nearly all genomes of the recently evolved Deuterostomia, while being much less common in the related Bilateria or other Metazoan lineages. Hence, the antiviral NTD of viperin-associated CMPK2 is a relatively recent innovation. Interestingly, most bird viperins lost the NTD but retained a mitochondria-targeting region (Supplementary Fig. 11). The N-terminal extensions found in other eukaryotes vary in length and have unknown functions, which warrant future investigations. Together these results demonstrate a functional genetic linkage that was not only maintained between viperins and CMPK2 kinases but incremented by the addition of the NTD in CMPK2 with an antiviral mechanism independent of viperin.

## Discussion

In this study, we investigated the evolutionary history of viperins across domains of life. Based on these results, we propose an evolutionary scenario, in which an ancestral substrate-promiscuous viperin was inherited from an archaeal ancestor by the First Eukaryotic Common Ancestor (FECA) (Fig. 4f, Fig. 5). We suggest that the initial repertoire of eukaryotic immune genes might have been primarily constrained by the available antiviral gene repertoire present in the ancestral Asgard archaea. This initial viperin then gave rise to a diversity of eukaryotic viperins through several mechanisms 1) serial innovations on the viperin gene itself, such as the emergence and selection of substrate specificity, and N-tail dedicated to localization to specific compartments in a eukaryotic cell; 2) Partnerships with genes maintained through genetic linkage, notably the associated kinase CMPK2.

**Figure 5:**
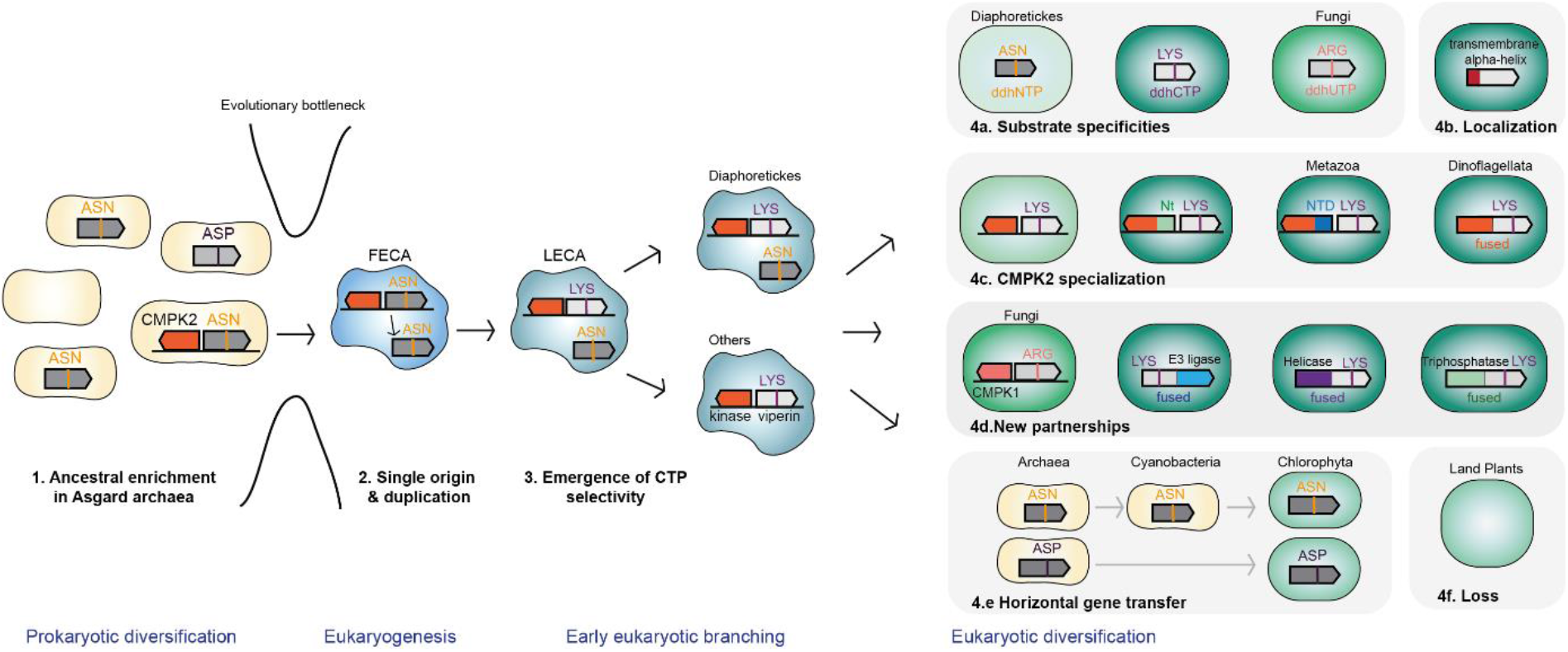
Proposed evolutionary scenario of viperin emergence and diversification in eukaryotes. **1**. The antiviral arsenal of Asgard archaea is enriched in viperins. **2**. The viperin gene was inherited during early eukaryogenesis, and has undergone duplication around eukaryogenesis, as suggested by the co-existence of the basal clade and distal clade in Diaphoretickes (Fig. 4f). **3**. One of the two variants underwent an extensive evolution that involved the development of a CTP-selective pocket along with other epistatic changes, while the other evolved slower and maintained the promiscuous pocket. Both variants remained in LECA, with the CTP-specific variant inherited by a great diversity of eukaryotic lineages, and the promiscuous version retained only by Diaphoretickes (Fig. 4f). **4**. Further diversification of viperins in eukaryotic lineages include **a**. the emergence of substrate selectivity; **b**,**c**,**d**. partnerships with additional functional domains through gene fusion and genetic linkage; **e**. HGT from prokaryotes; **f**. loss by specific eukaryotic lineages.

Viperins serve as a remarkable example of an immune gene that remains conserved in prokaryotes and eukaryotes. They have typically embraced adaptive strategies associated with genes that evolve slowly, such as precise specificity in biochemical interactions, forming partnerships with novel proteins, and maintaining functional genetic linkages. Similar evolutionary dynamics might be relevant for other eukaryotic immune systems that trace their origins back to prokaryotes. For instance, cGLRs, recently uncovered eukaryotic homologs of the human and prokaryotic immune gene cGAS^38,39^, also seem to show a more specific biochemistry in eukaryotes in the signal molecules produced as well as additional fused domains. However, contrary to viperins, cGLRs appear to have emerged from a relatively recent event of horizontal gene transfer^40^ and are thus restricted to a specific clade of eukaryotes. Whether similar evolutionary histories and dynamics are shared by other immune systems present in both prokaryotes and eukaryotes remains a topic that requires further elucidation.

The wide distribution and conservation of viperins among various organisms could be explained by their core antiviral mechanism, which targets a molecular process involved in the replication of many eukaryotic and prokaryotic viruses: RNA polymerization. Viperins appear to be more commonly found in eukaryotes compared to prokaryotes. This disparity could be attributed to fundamental differences in the viruses of prokaryotes and eukaryotes, as the eukaryotic virome is characterized by a majority of RNA viruses that heavily rely on RNA-dependent RNA polymerases (RdRps), while viruses relying on the host RNA polymerase dominate the prokaryotic virome^1,2^. Alternatively, potential additional antiviral mechanisms might be at play^18^. The loss of viperins in specific eukaryotic lineages, such as land plants, implies that these evolutionary dynamics are not universal among all eukaryotes. This divergence can lead to the selection of different immune mechanisms in different lineages.

We describe that viperins form and maintain functional partnerships with different genes through genetic linkage in eukaryotic genomes as exemplified by fusions and maintenance of genomic proximity of functionally associated genes like kinase CMPK2. Such genetic linkage seems to allow synergies to achieve an antiviral response. While functional genetic linkage has been heavily studied and described in prokaryotes, these occurrences are much more rare in eukaryotes^41,42^. Theoretical models predict the evolutionary conditions that could explain the emergence and maintenance of functional linkage in eukaryotic genomes^41,42^. The strength of the selective pressure imposed by viruses or the initial inheritance of an existing functional linkage could explain the maintenance of such gene neighborhoods. The extent to which this phenomenon occurs for immune genes, especially those inherited from prokaryotes, will require further investigation to ascertain its prevalence.

In this study, we demonstrate that the systematic search for eukaryotic counterparts of immune genes shared by both prokaryotes and selected eukaryotes offers a means to identify genes that uphold a consistent biochemical function within highly diverse eukaryotic genomes. These genes likely play a role in immune processes in these organisms. This approach offers promise in understanding immunity in organisms with limited knowledge of their immune systems. By revealing unique evolutionary markers specific to conserved immune genes, as seen with viperins and their associated kinases, we suggest these could serve as cues for discovering new immune genes in eukaryotes. Similar to how genetic linkage facilitated the discovery of numerous novel antiviral systems in prokaryotes^43^, this concept could be leveraged to unveil previously unknown immune genes in eukaryotic organisms.

## Materials and methods

### Detection of antiviral systems in archaea

Defense systems in archaea were detected using MacSyFinder v2.1.1^44^ using defense-finder-models v1.2.3^25^ and a minimal coverage of 0.4. The detection was made on The Genome Taxonomy Database (GTDB) v202 Archaea^45^. 50 Asgard archaea, or designated as Asgardarchaeota, genomes were excluded from the analysis as another larger dataset was used for Asgard archaea. Defense systems in Asgard archaea were detected using the same approach in our custom database of 496 in-house and publicly available Asgard archaea genomes (information and references provided in the Supplementary Table 7). Note that the majority of Asgard archaea and a fraction of the other Archaeal genomes are metagenome-assembled genomes, which can be incomplete and likely leads to a slight underestimation or contamination for the presence of defense systems.

### Construction of an HMM profile for viperins

To detect viperin homologs across prokaryotes and eukaryotes, we first constructed a new hidden Markov model (HMM) profile of viperin that encompasses its overall diversity. We first used known functional prokaryotic and human viperin sequences to construct the first HMM profile using HMMER v3.3.2^46^. We used this HMM profile to search across databases including GTDB v.207, EukProt v3, and our custom Asgard databases. The obtained sequences were used to construct a phylogenetic tree using IQtree v2.1.2^47^ model LG together with a set of known MoaA proteins as outgroups. MoaA proteins are distant homologs of viperin which do not show antiviral activity. Sequences branching with known viperin homologs are collected for HMM profile building, while sequences branching with MoaA were used as controls. To ensure an even sampling of diversity for HMM profile building, viperin sequences were clustered using CD-Hit v4.8.1^48^ with an identity threshold 0.65 (AA). The cluster representatives were aligned using MAFFT v7.49^49^ option L-INS-I, and the sequences with apparent N-terminal and C-terminal truncations were removed. This ultimately resulted in 140 well-aligned sequences, which we used to build a final viperin HMM profile. We searched the above viperin sequences and MoaA control sequences to set a cutoff for subsequent viperin searches.

### Detection of viperins

Two approaches were used to detect viperins depending on the objective: i) To describe the distribution of viperins in prokaryotic and eukaryotic genomes (Fig. 1C and Supplementary Figure 2): Based on the above viperin HMM profile, we searched for viperin homologs in GTDB v.207 (Asgard archaea excluded)^45^, a custom Asgard database (Supplementary Table 7), and EukProt v3^50^. These above three databases are used respectively for the statistical analyses of viperin presence in bacteria and archaea, Asgard archaea, and eukaryotes (Fig. 1c, Supplementary Figure 2). They are listed in Supplementary Table 2. In rare cases, a genome encodes more than 1 copy of viperins. ii) To explore the diversity of viperins: We used DefenseFinder^25^ to query a custom database comprising EukProt v3, GTDB v.202, GenBank from December 2021, and the custom Asgard database described previously to detect viperins (genomes listed in Supplementary Table 8). We added to these viperins prokaryotic viperins previously detected^20^. These viperins are listed in Supplementary Table 3 and used for structural analyses and detection of additional domains.

### Phylogenetic analyses of viperins

All detected viperins were first clustered using CD-Hit^48^ at a cutoff of 0.65 sequence identity. Previously examined bacterial viperins were used as preferred representatives of these clusters. These sequences were aligned using MAFFT^49^ using option auto, trimmed using TrimAl v1.4^51^ option gappyout. After removing some truncated sequences, the alignment was first phylogenetically analyzed using IQtree based on model LG. The tree was pruned using Treemmer v0.3^52^ to keep 207 sequences as backbone for the construction of three phylogenetic trees: 1) 207 viperins + 10 MoaA outgroup, which samples viperin diversity evenly across the kingdoms of life (Supplementary Fig. 2d); 2) 207 viperins + 57 additional eukaryotic sequences + 10 MoaA as outgroup, which expanded the main eukaryotic viperin branch (Fig. 1d); 3) 158 Archaea-eukaryote representatives selected from the above tree, to further confirm the close phylogenetic relation between Archaea and the basal eukaryotic clade (Supplementary Fig. 2e). All three trees were constructed as follows: viperin sequences were aligned using MAFFT option L-INS-I, trimmed using trimAl option gt 0.3, and phylogenetic analysis using IQtree using LG+C60+F+G4 mixture model^47^, with ultrafast bootstrap values calculated using UFBoot2+NNI from 2000 iterations. UFBoot2+NNI optimization was shown to avoid overestimation of branch support compared to other previous ultrafast bootstrap approaches^53^. The phylogenetic trees were visualized using iTOL^54^. Potential HGT events are indicated if multiple related eukaryotic viperins emerged within a prokaryotic branch, whereas singletons were considered potential sequencing contaminations.

### Strains and growth conditions

*E. coli* strains (Top 10, MG1655, MG1655 Keio ΔiscR) were grown in LB or LB agar at 37°C unless mentioned otherwise. The antibiotics ampicillin (100 μg/mL) and kanamycin (50 μg/mL) were added when appropriate.

### Plasmid construction

Viperin genes were codon-optimized and synthesized by Genscript Biotech Corporation with a TAA STOP codon. Each viperin gene was synthesized with flanking sequences for facilitating cloning upstream (ACCCGTTTTTTGGGCTAACAGGAGGAATTAACC) and downstream (GAATTCCCAGGCATCAAATAAAACGAAAGGCT). All plasmids were constructed using Gibson assembly in *E. coli* TOP10. Each gene was amplified using primers Vip-F and Vip-R, and cloned into the plasmid backbone pAB-pVip58^20^, linearized by PCR using primers Vector-F and Vector-R (primer sequences in Supplementary Table 9). The mutant viperin variants were constructed by PCR site-directed mutagenesis using the corresponding P1 and P2 primers in Supplementary Table 9, and cloned with Gibson assembly into the backbone pAB-pVip58 amplified with primers Vector-F and Vector-R. The sequences of viperins cloned and used in this study are listed in Supplementary Table 4.

### Plaque assays

*E. coli* MG1655 Keio ΔiscR strains transformed with viperin expression vectors were used for plaque assays as previously described^20^. Individual colonies of transformed strains were picked into 5 mL selective LB medium and incubated overnight with shaking at 37°C. To prepare plates, overnight cultures were diluted 1:100 in MMB agar (LB, 5 mM MgCl2, 5 mM CaCl2, 0.1 mM MnCl2, and 0.5% agar) supplemented with L-arabinose (final concentration 0.004%) for induction of Viperin expression. Serial dilutions of phage T7 lysate in MMB were dropped on top of the agar plates (4μL droplets) and dried. Plates were incubated at 37°C overnight.

### Structural analyses of the Viperin pocket

Alphafold structures of viperin homologs were computed using the ColabFold notebook.v.1.5.2^55^ The amino acid sequences used to generate the structures are listed on Supplementary Table 3. The crystal structures of the viperins from *M. musculus* bound to CTP (6Q2P), and *T. virens* bound to UTP (7N7I), were downloaded from the RCSB PDB database^56^. To perform the structural analysis of the nucleotide binding pocket, we aligned and superimposed all generated structures with the mouse viperin using the “super” command in PyMOL v2.4.0 (Schrödinger, LLC). Subsequently, to define the nucleotide binding pocket of all generated structures, we highlighted residues within a 4Å distance from the CTP substrate extracted from the mouse viperin crystal structure. This allowed us to manually identify the nucleobase-stabilizing residues at key positions within the β-8 loop of each homolog. We then constructed a structure-guided multi-sequence alignment using the PROMALS3D server^57^, with all the amino acid sequences in Supplementary Table 3 as input, and different substrate-bound viperin crystal structures as reference (PDB IDs: 6Q2P, 6Q2Q, 7N7H, 7N7I). This alignment was used to manually determine the stabilizing residues of homologs with nearly identical β-8 loop sequences.

### Preparation of cell lysates

*E. coli* MG1655 Keio ΔiscR transformed with viperin expression vectors, or expressing the negative control MoaA, were grown overnight in liquid cultures in selective LB medium. Overnight cultures were used to inoculate 50 mL of selective MMB medium (LB, 5 mM MgCl2, 5 mM CaCl2, 0.1 mM MnCl2) and incubated for 1h45 at 37°C with shaking (to reach an OD600 of 0.2-0.4). Gene expression was then induced with 0.2% L-arabinose, and cultures were further incubated for 90 minutes at 37°C with shaking. Bacterial pellets were recovered by centrifugation in Falcon tubes at 4°C. Samples were kept on ice throughout all following steps. Cell pellets were resuspended in 500μL cold methanol (50% v/v in ultrapure water, LC-MS grade), and transferred to 1.5mL Eppendorf tubes. Samples were flash frozen in liquid nitrogen and thawed on ice with frequent vortexing for 10 minutes. After repeating another freeze-thawing cycle, samples were centrifuged at max speed for 10 minutes at 4°C. The supernatants were transferred to Microcon®-5kDa centrifugal filters (Sigma-Aldrich) and centrifuged at max speed for 30 minutes at 4°C. The resulting flow throughs were transferred to glass vials and stored at -80°C until LC-MS analysis.

### Detection of ddh-nucleotides in cell lysates

LC-MS analysis was performed by MS-Omics (Vedbæk) as follows. Prior analysis, samples were evaporated under a gentle nitrogen stream. Extracts were reconstituted in an equal mixture of mobile phases A and B (140 μl, ammonium acetate in water: acetonitrile: medronic acid - 10 mM in 55: 44.9: 0.1 vol/vol/vol, adjusted to pH = 9) and filtered (0.22 μm). The samples were analyzed with a custom polar metabolite method, which is slightly modified from an existing protocol^58^. The analysis was carried out in a randomized order using a UPLC system (Vanquish, Thermo Fisher Scientific) coupled with a high-resolution quadrupole-orbitrap mass spectrometer (Q Exactive HF Hybrid Quadrupole-Orbitrap, Thermo Fisher Scientific). The ionization was achieved with an electrospray ionization interface operated in positive and negative ionization mode scanning a mass-to-charge range between 200 and 950. Peak areas were extracted using Compound Discoverer 3.0 (Thermo Fisher Scientific).

We used cell lysates from cultures expressing hVip and pVip58 as positive controls for the detection of masses matching derivatives of ddh-nucleotides previously reported^20^. Specifically, we focused on identifying the presence of ddhCTP through the detection of ddh-cytidine (ddhC, *m/z* 225.075) and ddh-cytidine monophosphate (ddhCMP, *m/z* 305.041) as markers. Additionally, the masses associated with ddhUTP (*m/z* 465.958), ddh-uridine diphosphate (ddhUDP, *m/z* 385.992), ddhGTP (*m/z* 504.980), and ddh-guanosine monophosphate (ddhGMP, *m/z* 345.047) were monitored to detect the synthesis of ddhUTP and ddhGTP, respectively. MS/MS data confirmed that the fragmentation spectra of these masses correspond to expected compounds (Supplementary Table 10).

### Detection of N-terminal tails of viperins

Viperins with a length between 250-400 amino acids and presenting both GGE and CxxxCxxC conserved motifs^23^ (involved in S-adenosylmethionine and iron-sulfur cluster binding respectively) were chosen to screen for the presence of an N-tail. We used the TMHMM webtool to detect the presence of N-terminal transmembrane alpha helices (https://services.healthtech.dtu.dk/services/TMHMM-2.0/). Such viperins were excluded from the following analysis. Remaining viperins were aligned using MAFFT.v7.505^49^ (default parameters). Homologs presenting more than 25 amino acids (minimum number of amino acids observed in viperins with N terminal tails) before the beginning of the conserved central domain were considered as having a non-alpha helix N-tail. Analysis results are listed in Supplementary Table 3.

### Detection and phylogenetic analyses of viperin-related nucleotide kinases

CMPK2 belongs to a larger superfamily of nucleoside monophosphate kinases (NMPKs), which also contains thymidylate kinases, adenylate kinase, and CMP/UMP kinases^59^. The two CMP/UMP kinases named CMPK1 and CMPK2 are phylogenetically distinct. We first used a group of known NMPKs from eukaryotes, archaea, and bacteria to build an HMM profile via HMMER as described for viperin above. Then the HMM profile was searched in the databases to obtain NMPK homologs. To carry about phylogenetic tree analysis, CD-HIT was used to remove the highly similar sequences (identity threshold 0.65). The rest of the sequences were aligned using MAFFT option auto, highly flexible regions were removed by TrimAl option gappyout, and built an initial large tree using FastTree v2.1.0 using default settings^60^. This initial tree was pruned using Treemmer to result in 1500 sequences. We re-aligned these remaining sequences using MAFFT option localpair and carried out maximum-likelihood analyses using IQtree using model LG+C10+F+G4+PMSF^61^. We then compared the genomic positions of NMPKs with the viperins to examine whether they constitute neighbors or gene fusions. Only genes having less than 3 genes in between were highlighted in the trees. The alignment of selected CMPK2 sequences were plotted with TeXshade v1.26^62^ to show the N-terminal extensions. The mitochondria-targeting peptides were predicted using TargetP 2.0^63^.

### Detection of additional protein domains

Fifty-three eukaryotic Viperins with a length above 500 amino acids were analyzed using the hmmsearch module of HH-Suite3^64^. All HMM profiles of the Pfam-A database were searched against the amino acids sequences of the selected Viperins, using the pre-defined GA gathering threshold of each profile of the Pfam database (--cut_ga option). The fifty-three viperins were also analyzed using HHpred (https://toolkit.tuebingen.mpg.de/tools/hhpred) to search for homology against HMM profiles of the COG-KOG database^65^, with minimal probability of hits set to 95%. Domains predicted as E3-ubiquitin ligase domain are KOG1812 or PF01485.22, KOG1814; domains predicted as predicted triphosphatase or cyclase domain are COG2114 or PF00211, COG1437.

## Data availability

Data used for this study are available here: https://github.com/mdmparis/viperins_evolution_2023

## Acknowledgements

We are grateful to members of the MDM lab, Eduardo Rocha, Tanita Wein and Benjamin Morehouse for their useful comments on earlier versions of the manuscript. Several bioinformatic analyses were performed on the Core Cluster of the Institut Français de Bioinformatique (IFB) (ANR-11-INBS-0013).

## Contributions

Y.F. and F.T. performed computational analyses for the detection of anti-phage defense systems in archaeal genomes. Y.F., J.C., F.T., H.S., H.G. and F.W. performed computational analyses related to viperin detection and phylogeny. H.S. and H.G. constructed strains and performed all experiments with assistance from B.O. H.S. analyzed mass spectrometry data, performed structural analyses and designed mutated viperins. H.S and H.G. performed computational analyses of N-terminal tails. H.G. performed analyses of viperin fused domains. Y.F. and J.C. performed computational analyses of viperin-associated nucleotide kinases. F.W. and A.B. supervised the project. H.S., H.G., F.W. and A.B. wrote the manuscript. All authors contributed to the review of the manuscript and provided final approval of the work.

