## Supplementary Materials for "Viperin immunity evolved across the tree of life through serial innovations on a conserved scaffold"

### Supplementary Tables

**Supplementary Table 1 -** DefenseFinder analysis results
**Supplementary Table 2 -** Detected viperin genes in the genomes of bacteria, archaea, and eukaryotes.
**Supplementary Table 3 -** Sequence and structural analysis of detected viperins
**Supplementary Table 4 -** Viperins tested *in vivo*
**Supplementary Table 5 -** Viperin-neighboring kinases
**Supplementary Table 6 -** List of viperins with detected additional domains
**Supplementary Table 7 -** Asgard genomes used in this study
**Supplementary Table 8 -** Genomes used to detect viperins diversity
**Supplementary Table 9 -** Primers used in this study
**Supplementary Table 10 -** MS-MS spectra of detected ddh-nucleotides

### Supplementary Figures

**Supplementary Figure 1 -** Defense systems in archaeal genomes
**Supplementary Figure 2 -** Maximum likelihood phylogenetic analyses of viperins
**Supplementary Figure 3 -** Conservation of viperin's active site
**Supplementary Figure 4 -** Antiviral activity of all viperin homologs experimentally tested
**Supplementary Figure 5 -** Detection of ddh-nucleotides in viperin-expressing cultures
**Supplementary Figure 6 -** CTP and UTP selective pocket variants
**Supplementary Figure 7 -** ASP pocket variants
**Supplementary Figure 8 -** Viperin N-tails
**Supplementary Figure 9 -** Genomic neighborhoods of viperin-kinase gene pairs
**Supplementary Figure 10 -** Maximum likelihood analysis of CMPK2
**Supplementary Figure 11 -** CMPK2 sequence alignment

### Supplementary Figures

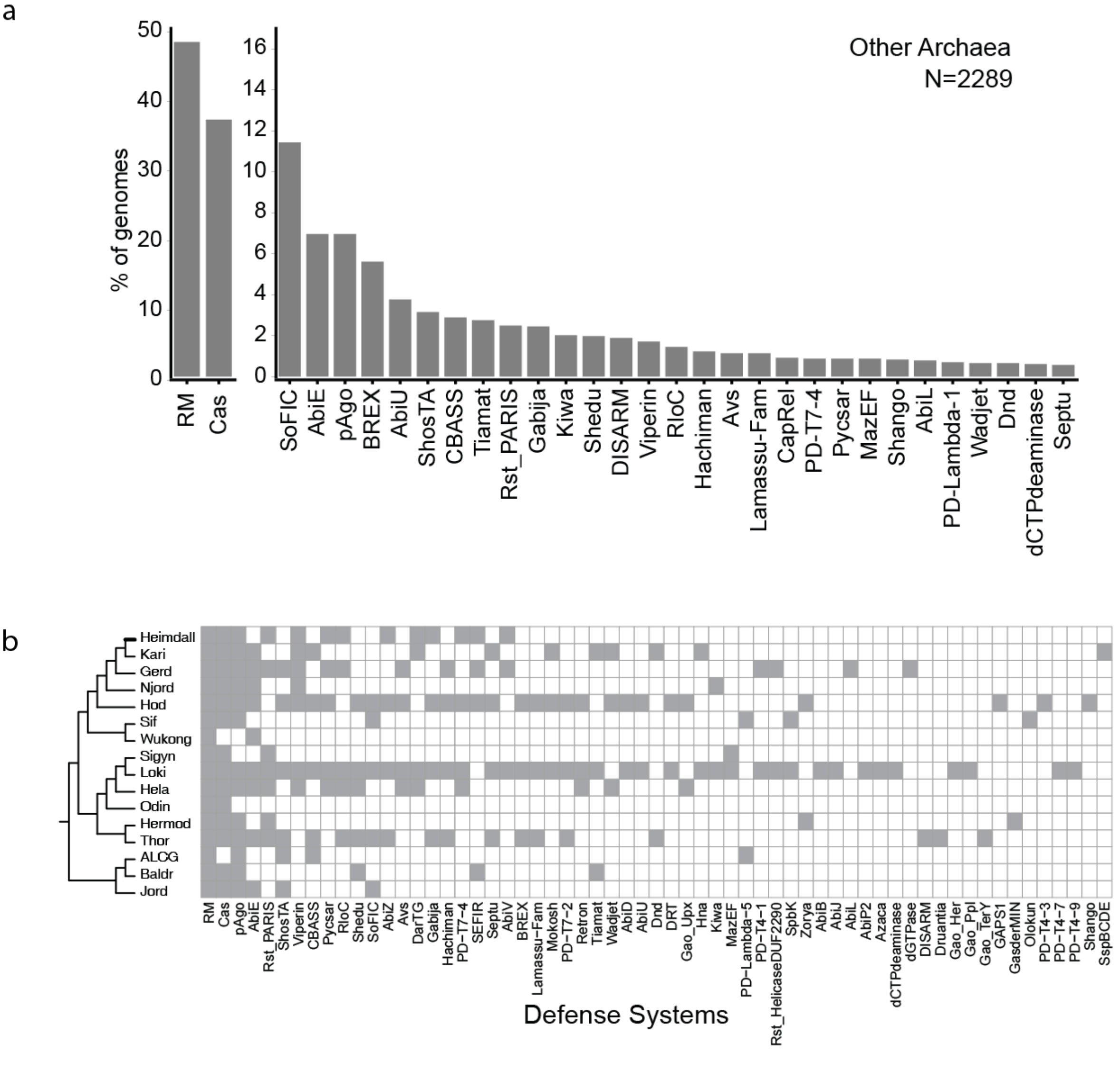


**Supplementary Figure 1. a.** Detection of known prokaryotic defense systems across diverse archaea (excluding Asgard archaea), ordered based on the percentage of analyzed genomes that encode them. **b.** Distribution of defense systems in Asgard archaeal genomes. Distribution exhibits varying levels of patchiness across different lineages of available Asgard archaea genomes as detected by DefenseFinder. Grey and blank represent presence and absence, respectively.

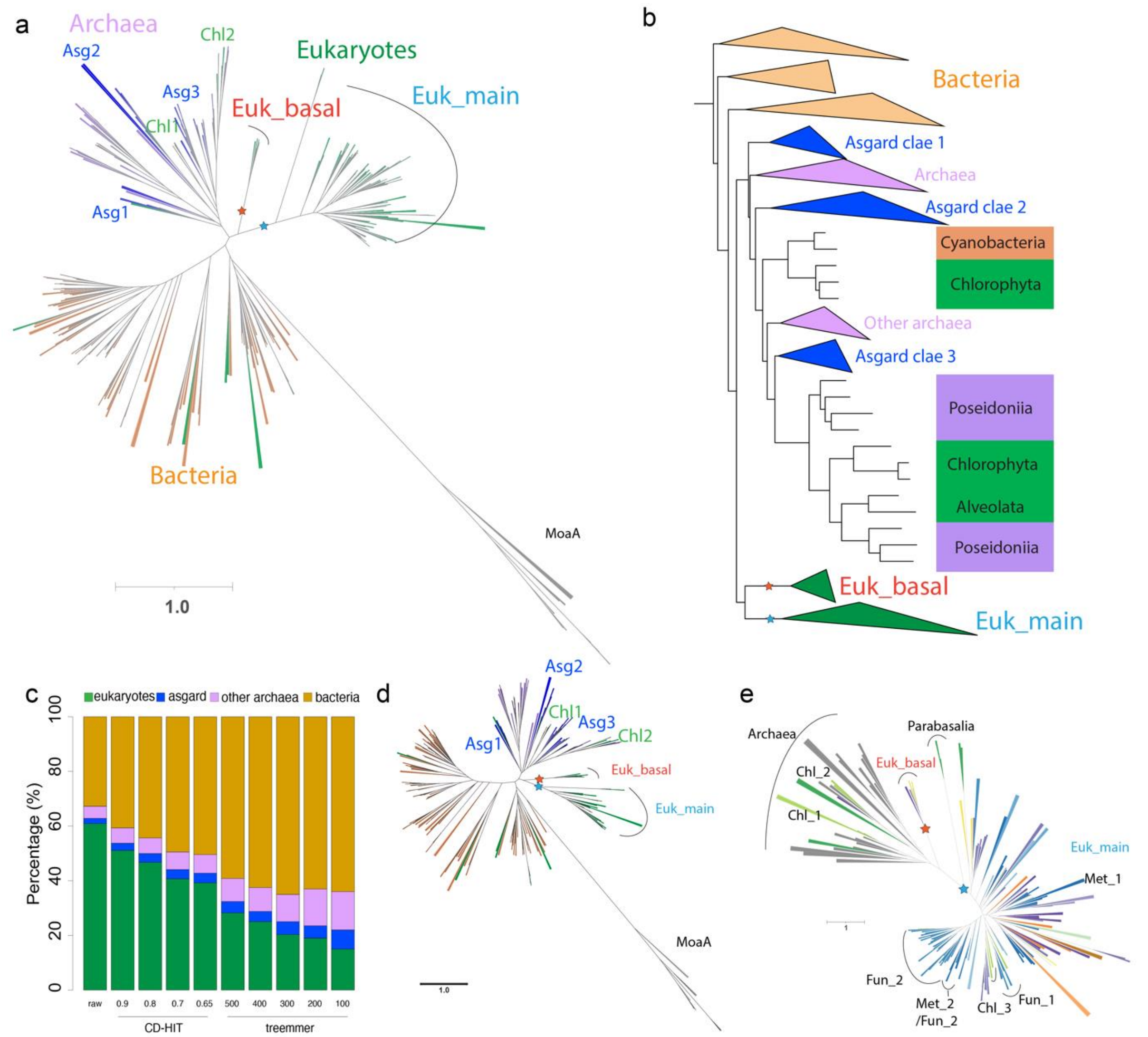


**Supplementary Figure 2. Maximum likelihood phylogenetic analyses of viperins. a.** Unrooted tree using an expanded selection of eukaryotic viperins (same as Fig. 1d). Asgard archaeal viperins diverged early into 3 distinct branches, indicated as Asg1, Asg2, and Asg3. Two Chlorophyta viperins originated from within the archaeal clade, indicated as Chl1 and Chl2. The basal and distal (main) eukaryotic clades are indicated by stars in red and blue, respectively. **b.** Collapsed tree (same as Fig. 1d) highlighting potential horizontal gene transfers: 1) archaeal viperins into cyanobacteria and subsequently into Chlorophyta, and 2) archaeal viperins into Chlorophyta and Alveolata. **c.** Sequence similarity and phylogenetic distance based down-sampling of viperins demonstrate the diversity of viperins detected in this study. It shows that while eukaryotic viperins constitute the majority of the dataset, they contain highly similar sequences. Detected bacterial viperins have the highest phylogenetic diversity. The fractional abundance of Asgard archaeal and other archaeal viperins increased most dramatically as closely related viperins are clustered, indicating that they are undersampled in the current genomic datasets. Future discoveries of more archaeal genomes may provide more insights into the evolutionary transition from archaeal viperins to eukaryotic viperins. CD-HIT approach downsampled sequences based on identity cutoffs (from 0.9 down to 0.65); Treemmer approach sets the number of retained sequences after tree pruning (from 500 down to 100). **d.** Phylogenetic analyses of viperins with reduced number viperins, which are relatively evenly sampled with respect to their phylogenetic distances, showing the same global topology as 1a. **e.** Phylogenetic analyses of diverse eukaryotic viperins showing the close phylogenetic distance between the basal eukaryotic viperin clade, and its apparent phylogenetic distance with the main viperin clade that is distal to archaeal viperins. Branch colors are identical to the eukaryotic tree of life depicted in Fig. 4f.

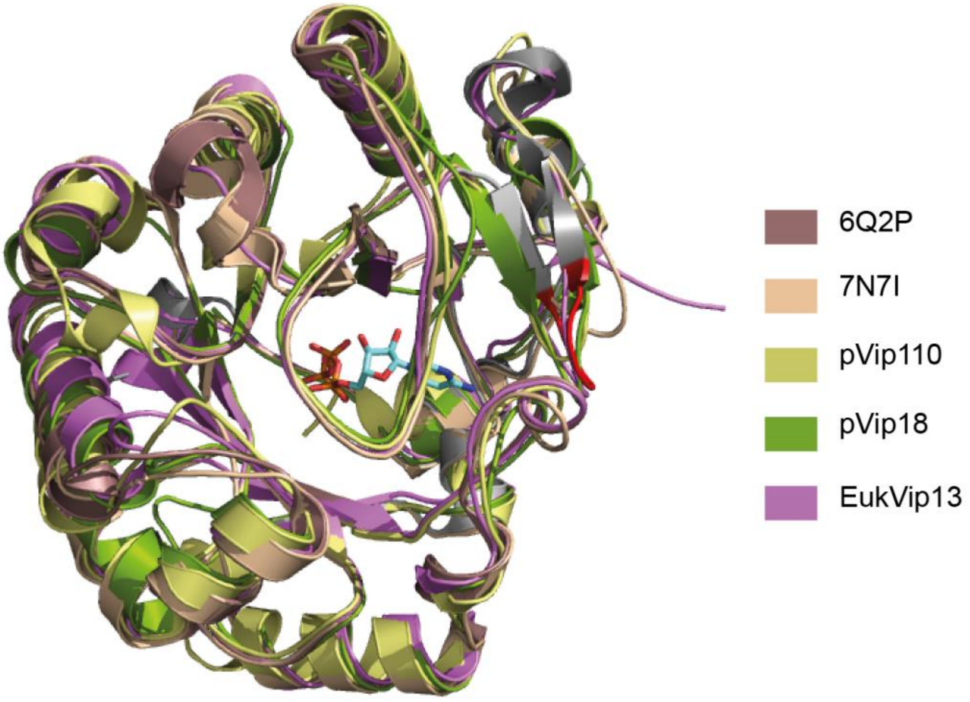


**Supplementary Figure 3: Conservation of viperin's active site.** Aligned structures of crystallized viperin from *M. musculus* (PDB ID: 6Q2P), *T. virens* (PDB ID: 7N7I), and Alphafold models of pVip110, pVip18 and EukVip13, demonstrating the conservation of the barrel fold that contains the active site. Highlighted in red is the flexible β-8 loop of the mouse viperin that contains residues involved in nucleobase binding. CTP is represented as sticks.

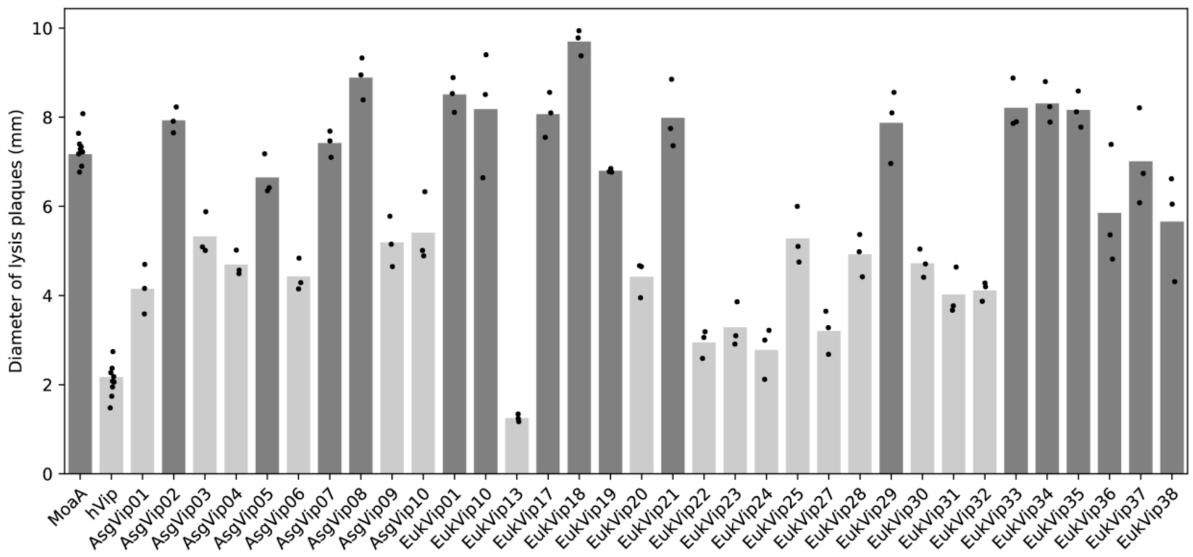


**Supplementary Figure 4**: **Antiviral activity of all viperin homologs experimentally tested**. Diameter of phage T7 lysis plaques infecting *E. coli* expressing viperins or the control MoaA. Bar graphs represent an average of three biological replicates, with individual data points overlaid. Light gray bars display viperins that display statistically significant differences compared to the MoaA control (one-sided t-test corrected by Bonferroni, p-value<0.01); these homologs are considered to have antiviral activity. Data for the tested viperin EukVip26 are not displayed since its expression was toxic in *E. coli*.

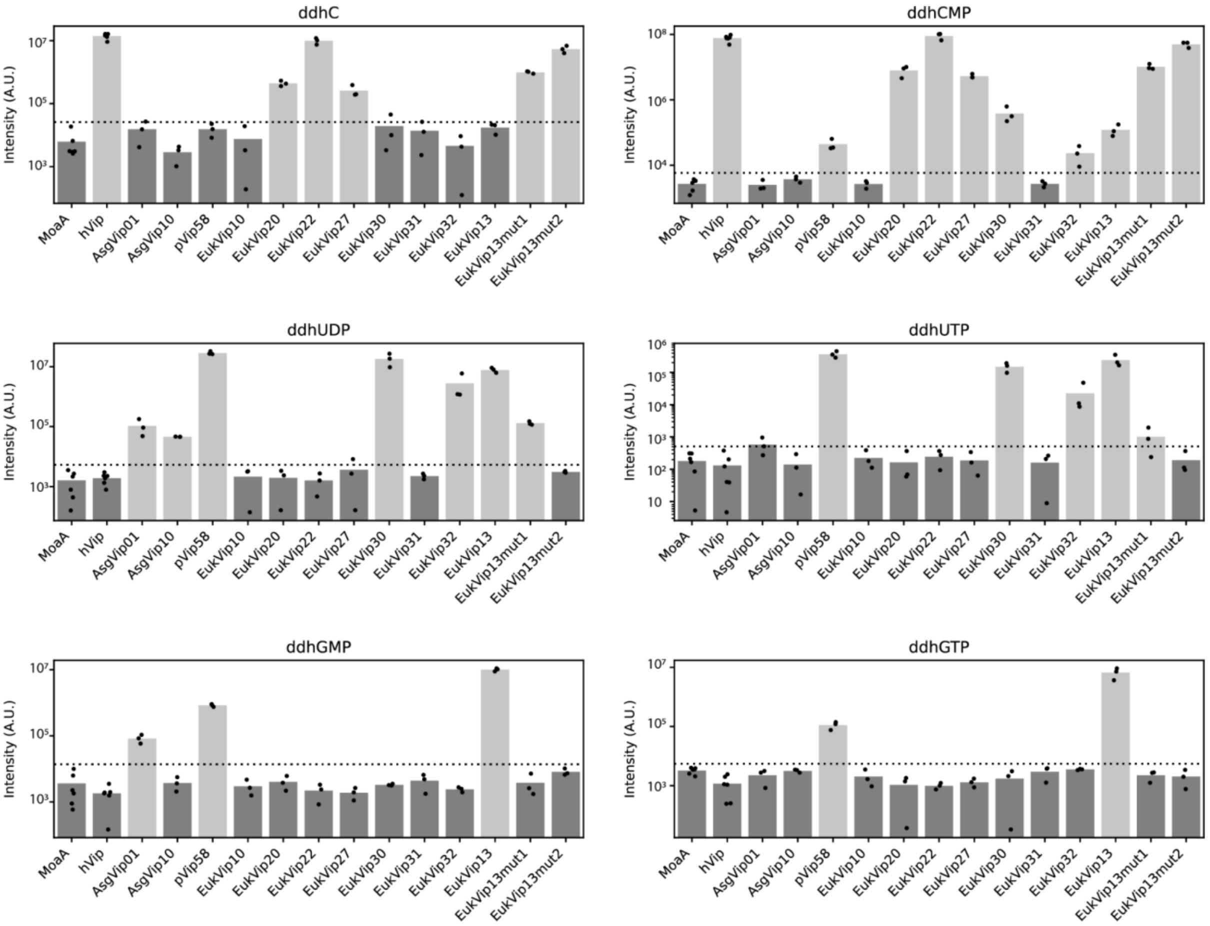


**Supplementary Figure 5**: **Detection of ddh-nucleotides in viperin-expressing cultures.**
Bars represent the average of biological replicates with individual points overlaid. Horizontal dotted lines correspond to the limit of detection (LOD) for each compound. Bars in light gray highlight samples with compound signals above the LOD.

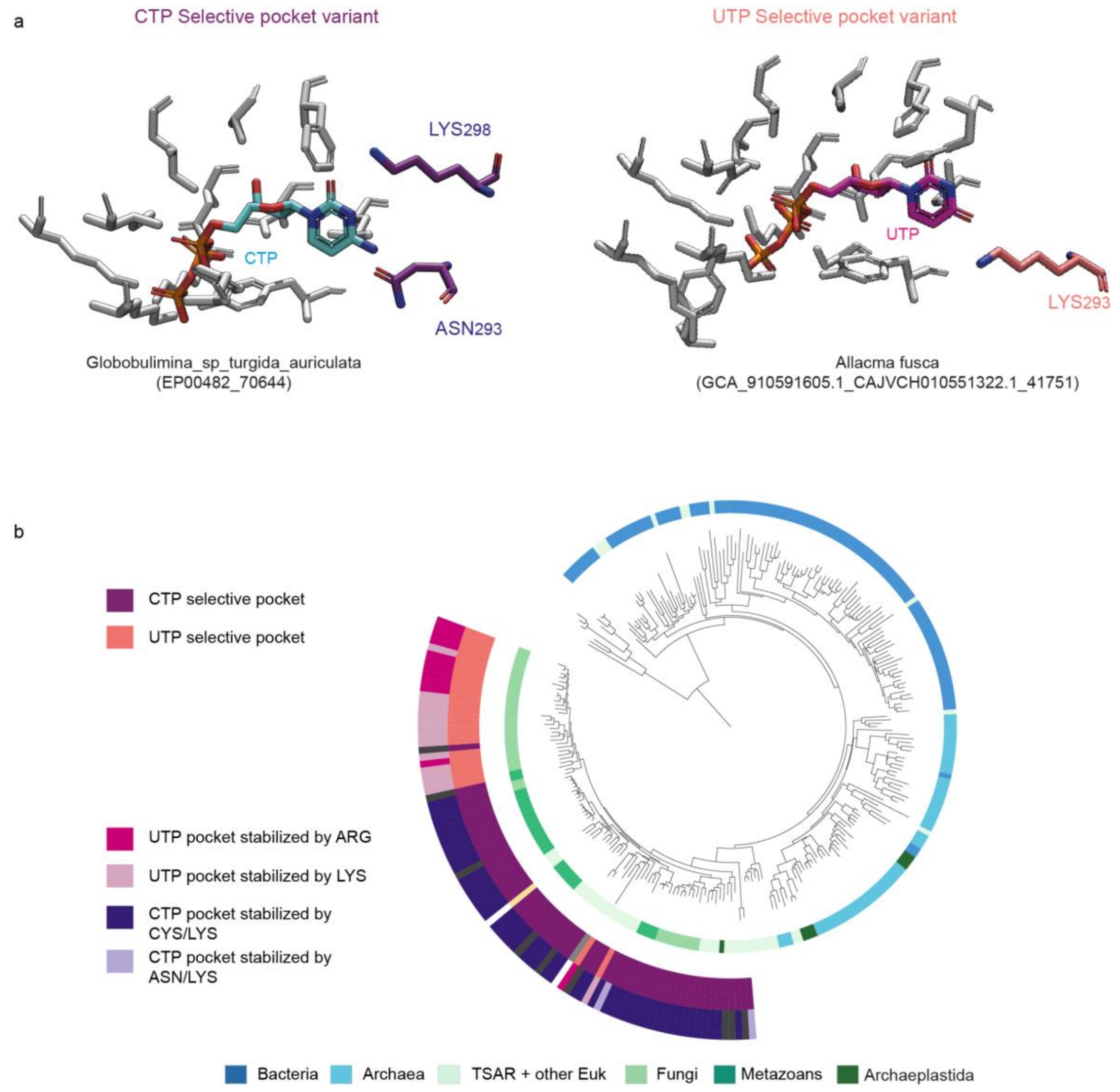


**Supplementary Figure 6**: **CTP and UTP selective pocket variants. a.** Magnified cutaway of additional variants of the selective nucleotide binding pockets in viperins structures (AlphaFold models). Putative CTP selective pockets were observed with an asparagine residue at the position of Cys314 in the viperin of *M. musculus*, as exemplified on the left. Putative CTP selective pockets were observed with an asparagine residue at the position of Cys314 in the viperin of *M. musculus*, as exemplified on the left. Putative UTP selective pockets with a lysine residue (at the position of Arg257 in the viperin from *T. virens*) that would form specific H-bonding interactions with the uridine base of UTP. **b.** Distribution of the different selective pocket variants on the viperin tree.

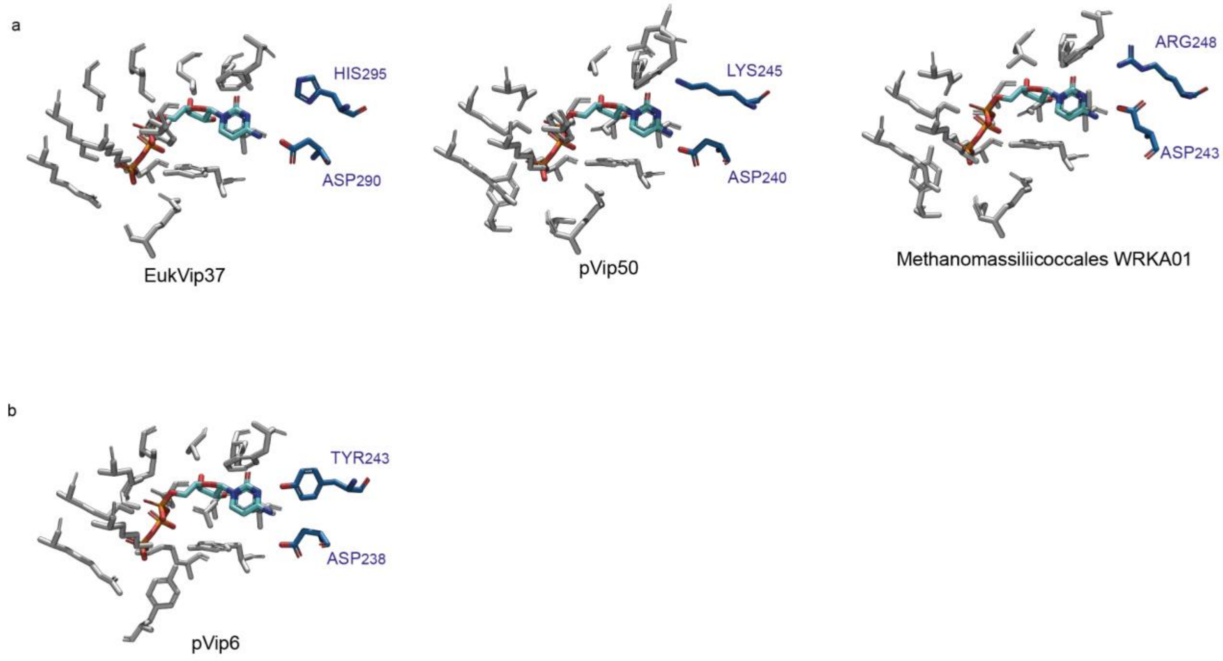


**Supplementary Figure 7**: **ASP pocket variants.** These pockets are characterized by a conserved aspartic acid residue at the position of Cys314, and diverse residues at the position of Lys319 of the mouse viperin: either (**a**) a positively charged residue or (**b**) a tyrosine . Previous studies show that viperins with these pockets, can selectively utilize CTP (pVip3, pVip50 and pVip6 - with a lysine or tyrosine analogous to Lys319), and in some cases multiple substrates (pVip44 - with a histidine analogous to Lys319, (see Bernheim et al., 2021). More research is required to fully assess the substrate selectivity of these rare homologs.

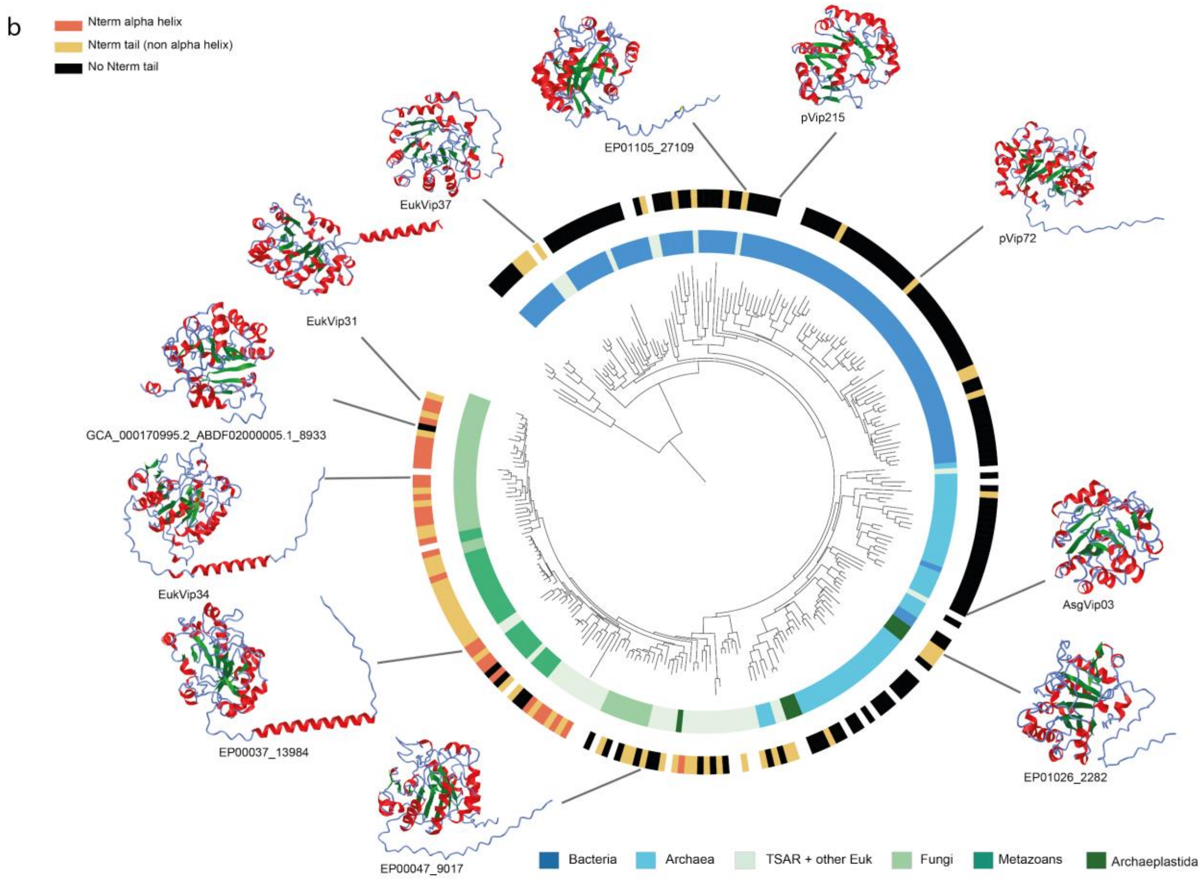


**Supplementary Figure 8: Viperins N-tails**

Representatives predicted structures of viperins with different N-tail configurations. Structures were predicted using ESMfold, plotted using iCn3D Structure Viewer (https://www.ncbi.nlm.nih.gov/Structure/icn3d/full.html, with alpha helices in red, and beta sheets in green), and mapped on the phylogenetic tree of viperins (from Fig 1.d). Outer ring corresponds to different N-tail configurations. Black indicates the absence of Ntail, yellow indicates a predicted N-terminal alpha helix, and orange indicates non alpha helix N-tail.

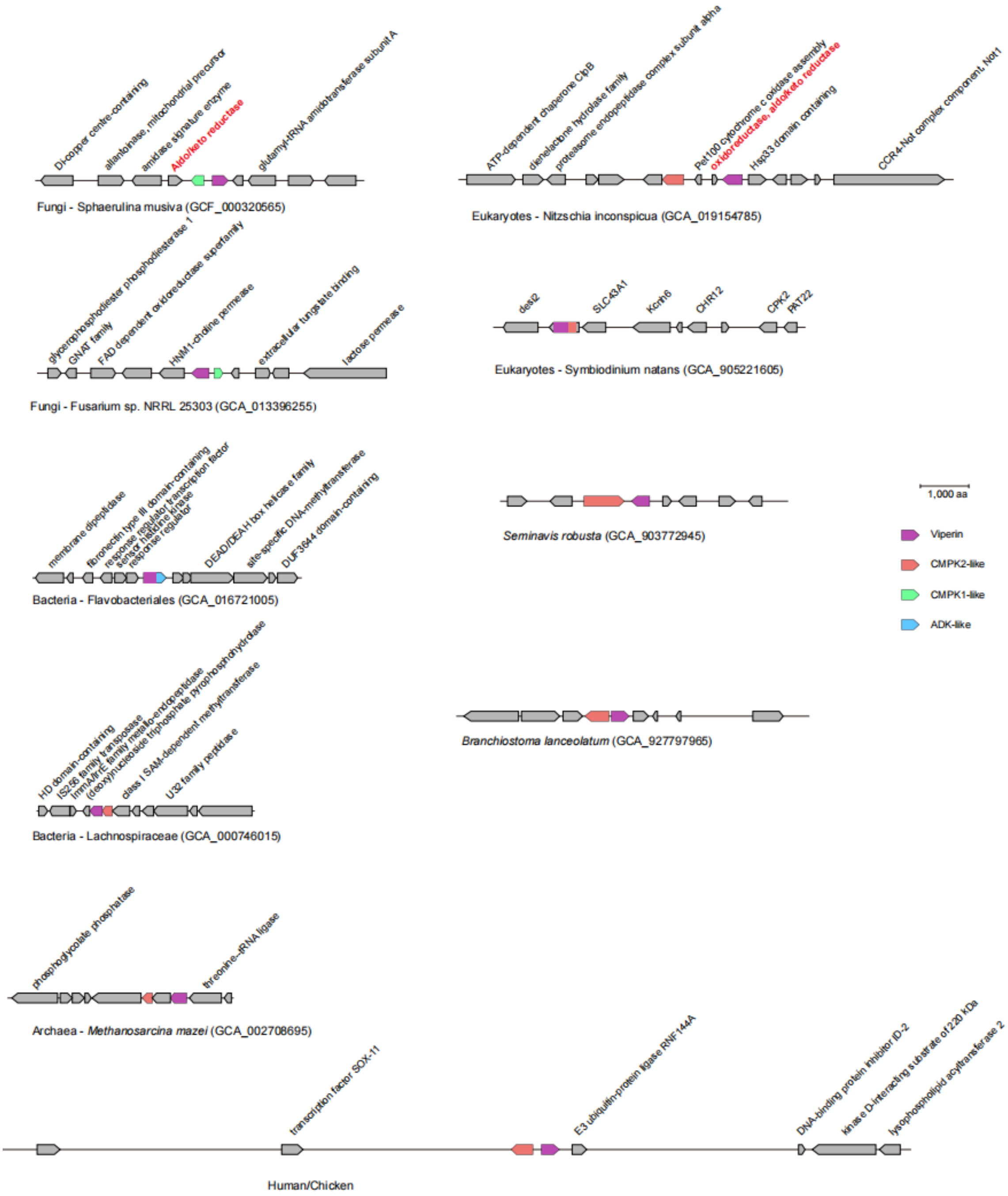


**Supplementary Figure 9. Genomic neighborhoods of viperin-kinase gene pairs.** The genes were scaled down by their amino acid length (introns not shown) for representation. Their relative distances on the chromosome were indicated. Genes with predicted functions were indicated.

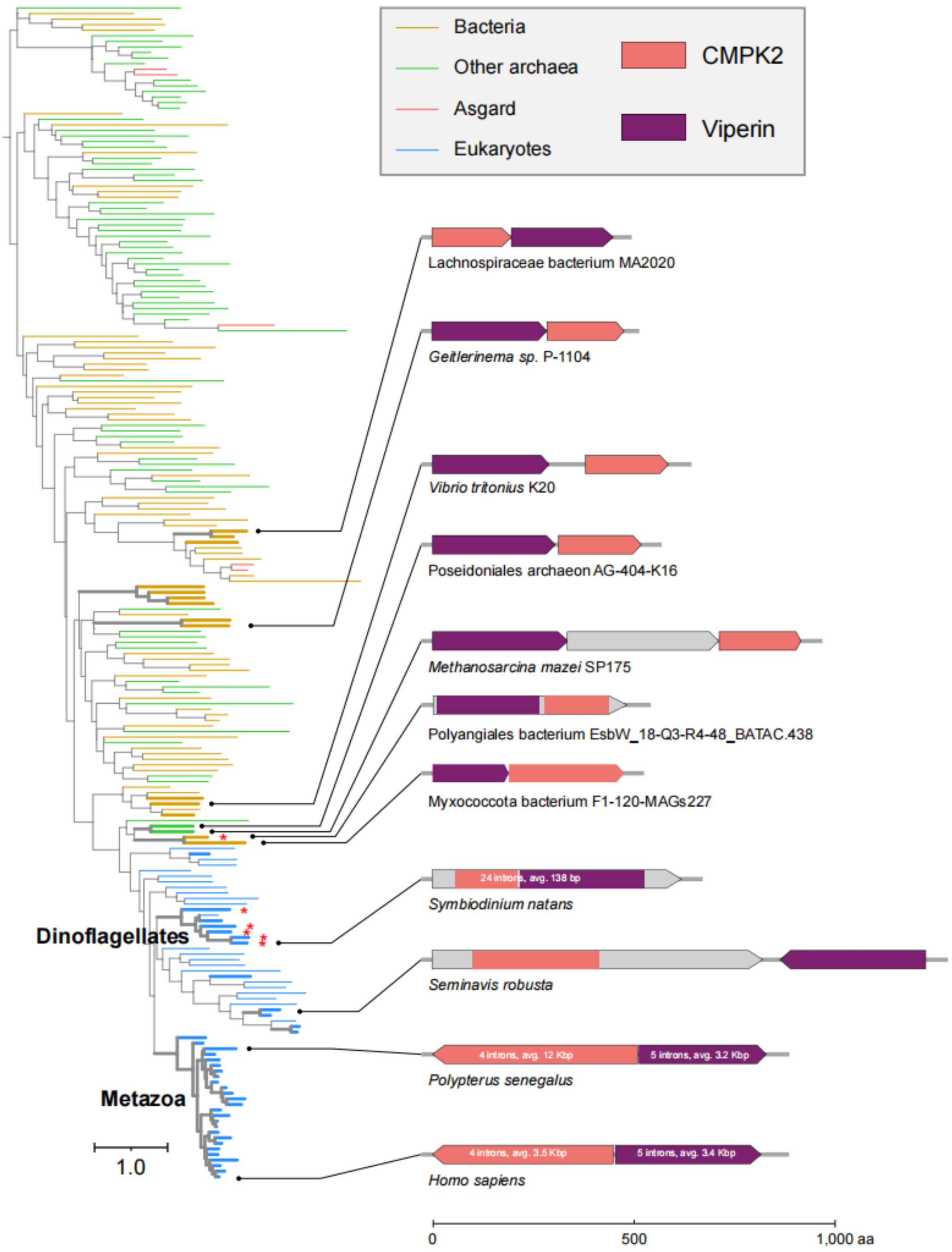


**Supplementary Figure 10. Maximum likelihood analysis of CMPK2 clade.** The thick branches on the tree show those close to viperin (within 3 genes). The red asterisks indicated CMPK2 and viperin fusion. Examples of genomic arrangements are shown on the right side. Genes were plotted based on their amino acid (aa) length, and their intron information was written on the gene wherever available.

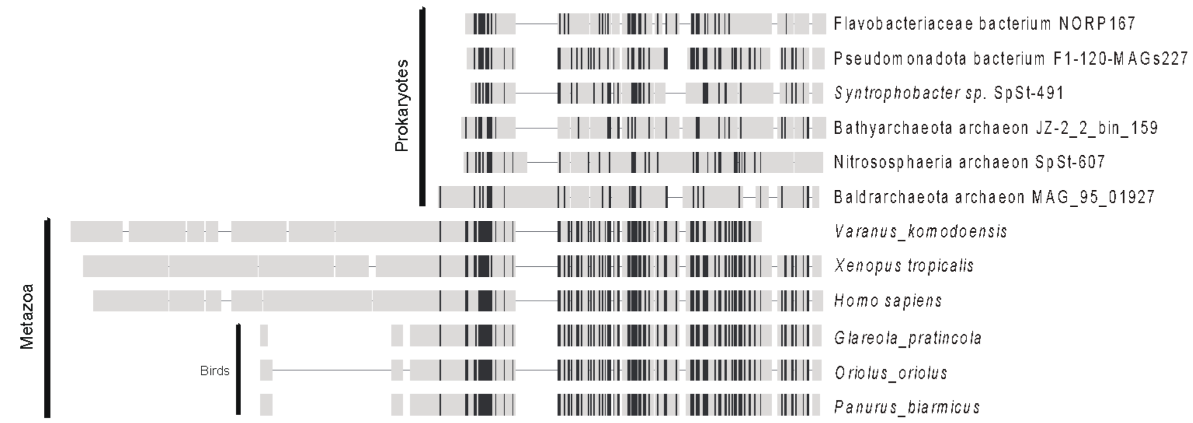


**Supplementary Figure 11. CMPK2 sequence alignment shows that the N-terminal domain in birds is much shorter than other metazoans.** TargetP predictions suggest that these shorter N-terminal sequences encode the mitochondria-targeting peptide. The black and gray colors show amino acids higher and lower than 50% identity, respectively.
